# Impaired lipoprotein secretion by APOE4 leads to lysosomal and mitochondrial dysfunction in human microglia

**DOI:** 10.64898/2026.05.12.724612

**Authors:** Jasmin S. Revanna, Karl Wessendorf-Rodriguez, Qiang Xiao, Thais S. Sabedot, Michael S. Cuoco, Snigdha Sarkar, Lucia Zhou-Yang, Christina K. Lim, Victoria N. Prozapas, Rowan S. Wooldridge, Jean Paul Chadarevian, Joshua M. Pratt, Sheila C. Steiner, Ava Katz, Jerome Mertens, Jeffery W. Kelly, Santiago Solé-Domènech, John T. Melchior, Christian M. Metallo, Jeffrey R. Jones, Fred H. Gage

## Abstract

While Apolipoprotein E4 (APOE4) is the greatest known genetic risk factor for late-onset Alzheimer’s disease, its mechanistic role in the brain-resident macrophage, microglia, remains elusive. Microglia are important in the clearance of pathology in disease, heavily relying on lysosome functionality; therefore, we sought to understand the impact of APOE4 on microglial function. APOE44 microglia have been shown to have lipid accumulation, yet the mechanisms leading to this accumulation are unknown. Using induced pluripotent stem cell-derived microglia, we found that the APOE4 haplotype resulted in transcriptional state shifts in microglia, suppressing activated-response microglia (ARMs) and promoting a G2 senescent-like state. We found that APOE44 microglia accumulate cholesterol esters and provide less lipid support to fibroblast-induced neurons, decreasing their synaptic connections. APOE44 microglia secrete significantly less lipoproteins, leading to the accumulation of lipoproteins within the cells including the lysosomes. APOE44 microglia exhibit impaired lysosomal acidification and degradation capacity. Further, our results elucidated that APOE44 microglia are proinflammatory and shift away from fatty acid oxidation towards glycolysis, due to dysfunctional mitochondria. Taken together, our findings indicate that a loss-of-function in lipoprotein secretion drives intracellular lipid accumulation, including within lysosomes, ultimately disrupting the lysosome-endoplasmic reticulum-mitochondrial axis. This drives a proinflammatory and metabolically compromised microglial phenotype with impaired neuro-supportive functions.

**GRAPHICAL ABSTRACT:** 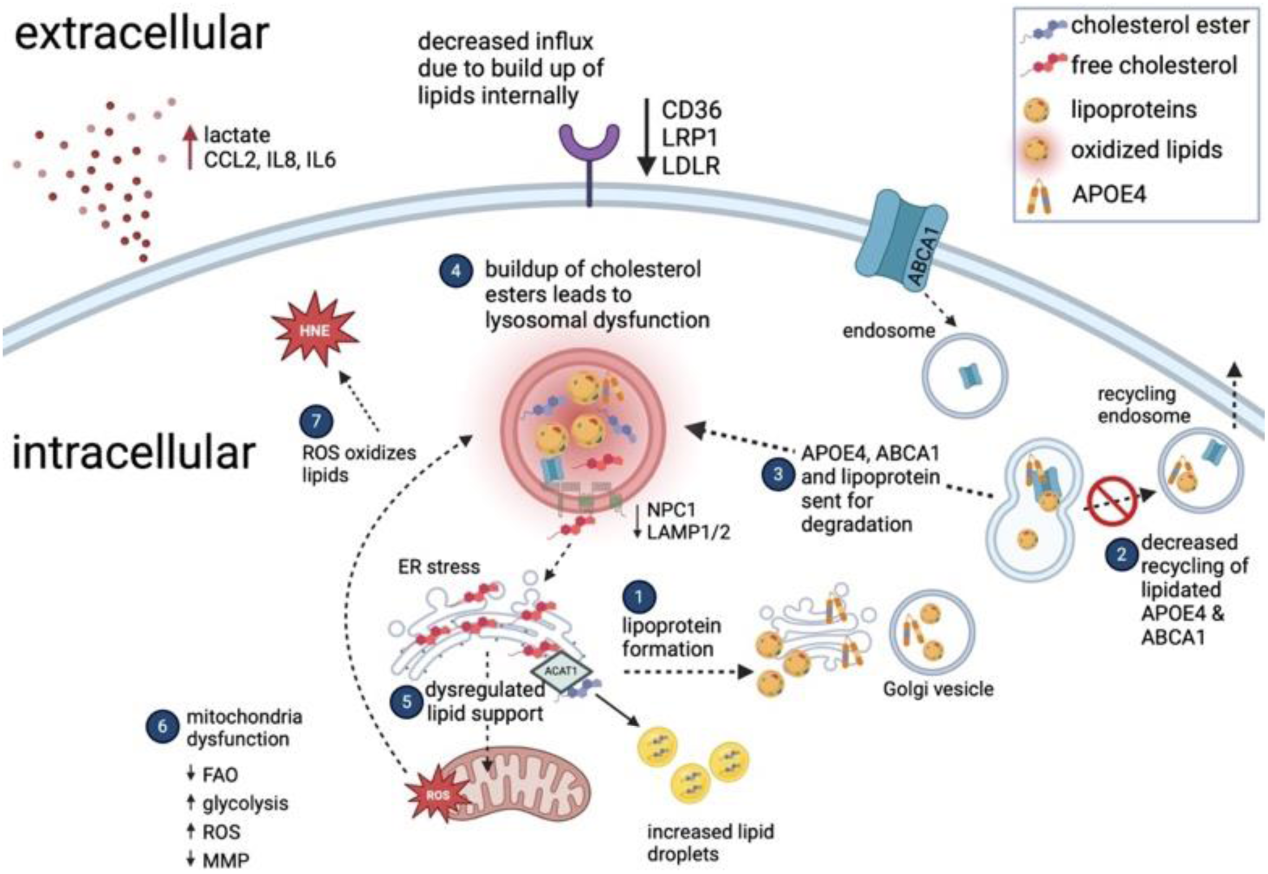

## INTRODUCTION

Microglia are specialized resident macrophages of the central nervous system that help to clear debris, pathogens, and neuronal synapses through phagocytosis^1^. Under pathological conditions, microglia can prune synapses excessively, increase pro-inflammatory cytokine release, and reduce trophic factor release^2,3,4^. The mechanistic role of microglia in the onset and progression of Alzheimer’s disease (AD) is not fully understood. The apolipoprotein E (APOE) haplotype is the major genetic risk factor for the development of AD, with the ε4 allele correlated to highest risk and the ε2 allele to reduced risk^5^. A recent study of homozygote carriers for APOE4 showed that, by age 65, >90% of APOE4 homozygote participants had abnormal levels of CSF_Aβ1–42_ compared to ∼30% of APOE33 carriers, highlighting a mechanistic difference in APOE4 carriers. In disease models, APOE4 expressing cells exhibit deposits of lipid droplets and changes in expression levels of cholesterol transport genes, yet how APOE4 contributes to this lipid accumulation remains unknown^6–11^. These findings emphasize the critical need to understand the role of APOE4 in the onset and progression of AD.

Microglia take on many states with distinct gene signatures, and the roles of each of these states are slowly being uncovered. Microglia surrounding amyloid plaques in genetic mouse models of AD have been characterized by their altered transcriptional profile and have historically been referred to as disease associated microglia (DAMs), also known as activated response microglia (ARMs)^12,13^. TREM2 knockout studies have shown that TREM2-deficient microglia lack ARMs and that this cluster is defined by clearance genes^14–16^. Microglia are the scavenger cells of the brain, where they clear debris through their active lysosomes^17^. A recent study identified that the knockout of BHLHE40/41 in human induced pluripotent stem cell (iPSC)-derived microglia (iMG) is sufficient to drive iMG into an ARM state, which the study referred to as DLAMs^14^. BHLHE40/41 knockout microglia exhibit an increase in lipids, lysosomal mass, degradative capacity and, importantly, are not proinflammatory^14^. It remains unclear whether this ARM population is beneficial or detrimental for disease and how the APOE4 haplotype may impact this state transition.

Emerging evidence indicates that microglia senesce in aged and AD mouse models^18–20^. Senescent cells secrete inflammatory cytokines [senescence-associated secretory phenotype (SASP)] that impact surrounding cells^21^. Recently, our lab showed that fibroblast-induced neurons (iNs) from AD individuals demonstrated increased populations of senescent neurons^22^. Microglial senescence is understudied, and it is unclear whether APOE4 microglia are more vulnerable to cell cycle arrest. Under healthy physiological conditions, microglia maintain a G0 quiescent state and proliferate when challenged^23–26^. This proliferation could be a catalyst for microglia to become senescent^18^. We sought to understand whether the APOE4 haplotype influences this transition into a senescent-like state through cell exhaustion.

To investigate the transcriptional state of microglia in APOE4 carriers, we used established methods^27^ to convert 22 iPSC lines from 5 APOE23, 6 APOE33, 7 APOE34, and 4 APOE44 individuals, and we performed single cell RNA sequencing (scRNAseq)^27^. In this *in vitro* model, we observed distinct transcriptional states mimicking what has been observed *in vivo*^12,14,15,28–33^. This approach allowed us to investigate how the APOE4 haplotype impacts microglial state shifts and how these shifts might impact onset and progression of disease. We demonstrated that APOE4 suppressed the ARMs, also known as DAMs, and promoted a G2 lipid-laden senescent-like state. This result suggests a neuroprotective role for ARMs. Isogenic APOE33 and APOE44 iMG revealed functional differences in the regulation of cholesterol esters, including an overall increase in cholesterol esters, and specifically their accumulation in lysosomes. Proteomics data reveal a downregulation of lysosomal hydrolases and V-ATPases in APOE44 iMG, which is supported by an impaired ability to acidify lysosomes upon challenge and reduced degradative capacity. Additionally, proteomics data indicate endoplasmic reticulum (ER) stress, which may dysregulate lipid support to the mitochondria. This dysregulation of lipid support to mitochondria is indicated by an increase in reactive oxygen species (ROS) and a decrease in mitochondrial membrane potential. Microglia can shift between utilizing glycolysis, oxidative phosphorylation, and fatty acid oxidation (FAO) to generate energy, depending on their activation state, with proinflammatory microglia mostly utilizing glycolysis and anti-inflammatory microglia utilizing FAO and oxidative phosphorylation^34^.

We found that APOE4 microglia contributed to a shift away from FAO towards glycolysis and were more proinflammatory. We were able to partially rescue these phenotypes with Liver X receptor (LXR) agonist GW3965. We show that iNs cultured with APOE44 iMG have significantly fewer lipids compared to those cultured with APOE33 iMG, reducing their synaptic connections, suggesting that APOE44 iMG may not provide neurons with appropriate lipid support. Lastly, APOE44 iMG secrete significantly less APOE and high-density lipoproteins (HDL) compared to APOE33 iMG. These data, taken together, indicate that microglia produce and export lipoproteins during normal function and that lipoprotein secretion is impaired in APOE44, resulting in a buildup of cholesterol esters and other lipids within the cells. These findings provide direct evidence for the role of APOE in microglia function in health and disease. Importantly, our results advance our understanding of the increased risk of APOE4 homozygosity and the onset and progression of AD.

## RESULTS

### Two-dimensional-induced microglia recapitulate in vivo mouse and human microglia transcriptional states

iPSCs from 14 AD and 8 healthy aged donors representing different APOE haplotypes (APOE23 n= 5, APOE33 n= 6, APOE34 n= 7, APOE44 n= 4; Table S1) were differentiated into iMG following established methods^27^. iMG were validated by a 6-marker microglial panel and FACS sorted (Figures 1A, S1A). This protocol includes first converting iPSCs into induced hematopoietic progenitor cells (iHPCs) and then into iMG (Figure S1B). Microglia are known to have many different states with different roles *in vivo* and even *in vitro*^12,14,15,28–33^. To characterize microglial states and the influence of the APOE haplotype, we performed scRNAseq. We captured high-quality whole cell transcriptomes from 40,723 iMG, with comparable numbers of cells from each line (Table S1, Figure S1C). Nearly all microglia express the microglial canonical markers AIF1, CST3, and CSF1R (Figure S1D). UMAP dimensionality reduction identified 9 distinct unbiased clusters that were annotated by creating module scores from previously identified clusters in other microglia studies (Figure 1B, 1C, Table S2)^14,15,28,29,31,33,35,36^. The resulting clusters were Activated Response (ARM), Ribosomal Response (RR), MHC Class II response (HLA(1) and HLA(2)), Homeostatic (HM), Female-enriched disease-associated microglia (FDAMic), G2 cell cycle, G1 cell cycle and Cytokine Response (CRM) (Figure 1B, Table S2). iMG derived from each iPSC line were well represented in each of the nine clusters (Figure S1E), and each cluster expressed canonical microglia markers (Figure S1F). Importantly, the ARM cluster overlapped well with both human and mouse ARM, DLAM and DAM signatures (Figure 1C, Table S2); henceforth, we refer to this cluster as ARM. This cluster expressed APOE, TREM2, FUCA1, C1QA, CD68 (Figure 1D, Table S2), lysosomal hydrolases (Figure S1G), and lysosomal V-ATPases ATP6V0C, ATP6V1F, ATP6V0E1 (Table S2), highlighting the cluster’s role in clearance^14,29,30,35^. Notably, this cluster had low expression of proinflammatory cytokines (CCL4, IL1B) and high expression of the homeostatic microglia marker MERTK (Figure S1H/I). Importantly, the ARM population had reduced expression of BHLHE40/41, whose KO in previous studies identified these transcription factors’ role in shifting microglia into an ARM state (Figure S1J)^14^. When analyzing each cluster by sex, one cluster was overrepresented by the female lines and overlapped with the signature of FDAMics (RPL13, RPLP1, SPP1, RPS19, ACTB, RASGEF1B, PSAP, CSF1R) (Figure S1K/L)^36^. Comparison of female and male microglia revealed higher expression of TLR7 and TLR8 in females (Figure S1M). Both genes are located on the X chromosome and have been reported to escape X-inactivation in other immune cell types^37^. Our data suggest that TLR7 and TLR8 may similarly escape X-inactivation in microglia, potentially contributing to an enhanced immune responsiveness in female cells.

**Figure 1.**
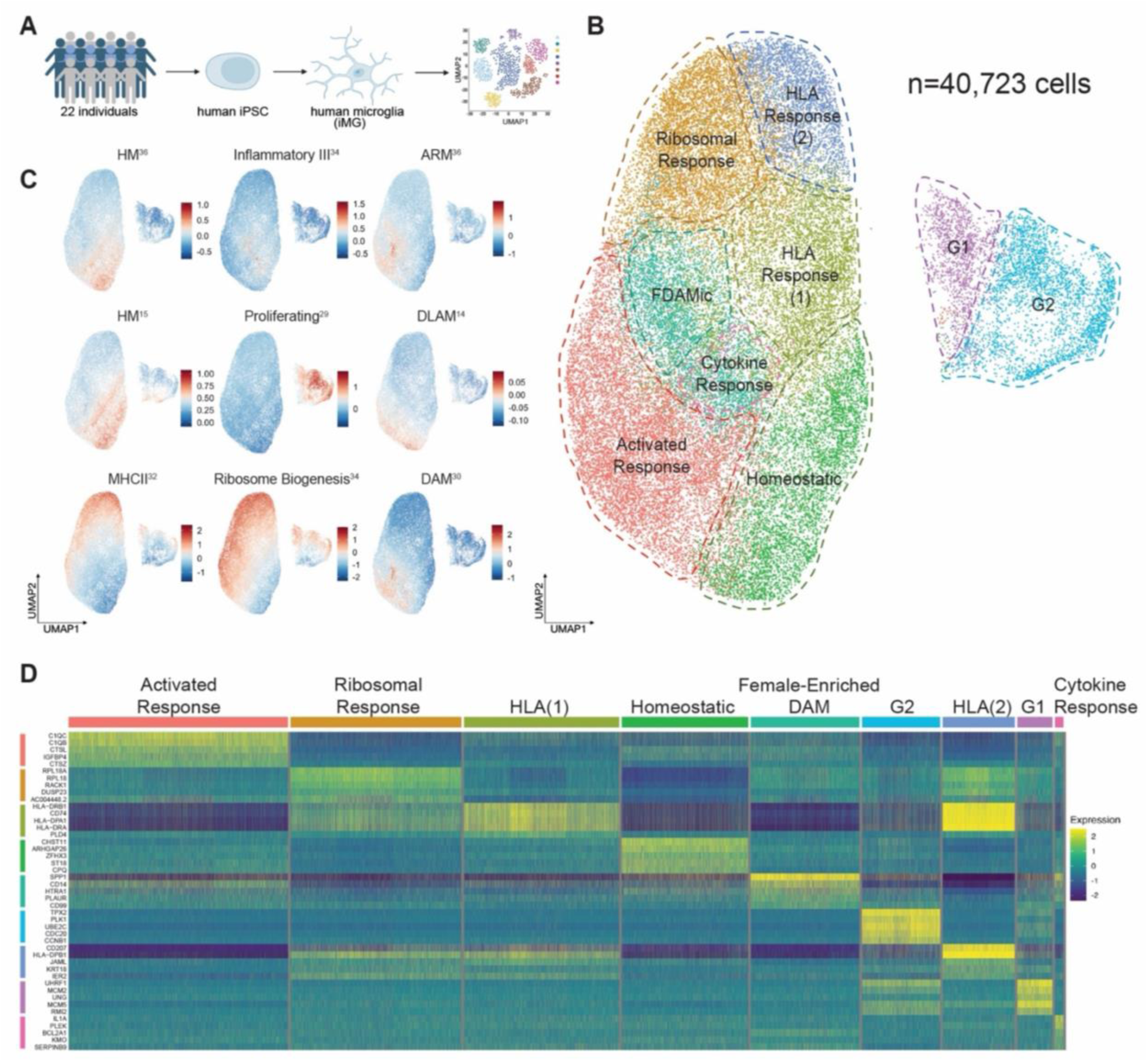
Two-dimensional-induced microglia recapitulate in vivo mouse and human microglia transcriptional states. (A) Schematic of donor cohort. (B) UMAP representation of all cells (n=40,723) from scRNAseq, with transcription states annotated. (C) Module scores of different transcriptional states from various microglia studies plotted onto our UMAP. (D) Heatmap of top 5 gene markers from each cluster shown (normalized expression scaled by gene is shown). See also Figure S1.

The ribosomal response cluster was defined by high expression of several ribosome-related genes, including RPL30, RPL19, RPL10, RPS8, and RPL18 (Figure 1D, Table S2), which may indicate that this population of cells is important in translation or is experiencing ER stress. The HLA(1) cluster is defined by MHC class II genes such as HLA-DRB1, HLA-DRB5, and HLA-DRA; furthermore, the HLA(2) cluster is defined by both ribosomal and MHC class II genes (Figures 1D, Table S2).

The homeostatic cluster is enriched in known homeostatic markers like PLXDC2, MERTK, and TGFBR1 (Figure 1D, Table S2)^38^. Our cytokine response microglia are defined by CCL4, CCL3, IL1B, NFKB1 and SOD2 and make up a small, but distinct, proportion of the entire dataset (Figures 1D, Table S2). Lastly, we have two clusters that express G1 cell cycle (MCM2, MCM3, MCM5) and G2 cell cycle (PLK1, TPX2, H2AFX) markers, which are thought to be proliferating microglia (Figures 1C, 1D, Table S2). Although our 2D iMGs recapitulate many transcriptional states observed *in vivo*, key homeostatic markers such as SALL1 and TMEM119 remain absent. Prior studies have demonstrated that microglial identity is strongly influenced by culture context, with notable differences between 2D systems and primary, 3D, or transplanted microglia^30,31,39,40^. Nevertheless, we leveraged the controlled, cell-autonomous nature of 2D culture to specifically interrogate how APOE haplotype influences microglial state transitions.

### APOE4 influences state shifts in microglia, suppressing the ARM signature

By separating the 22 donor lines by APOE haplotype (APOE23 n = 5, APOE33 n = 6, APOE34 n = 7, APOE44 n = 4), we sought to understand how the APOE4 haplotype may suppress or promote specific states in the scRNAseq dataset (Figure 2A). We asked if microglia from each haplotype are enriched in any specific cluster, and we visualized the distribution of each haplotype in the UMAP space (Figure 2B). Interestingly, we saw a significant increase of APOE4 iMG represented in the FDAMic cluster, largely driven by the female APOE4 iMG (Figures S2A-B), and an increasing trend in the ribosomal response cluster and the G2 cell cycle cluster (Figures 2B-C). The ‘proliferating’ clusters in a majority of microglia studies have been discarded and thus represent an understudied population. We saw a significant decrease in APOE4 iMG represented in the ARM cluster (Figures 2B-C, 2S2A). To understand if a reduction of ARMs correlated to disease prevalence, we compared single nuclei RNAseq datasets generated from human postmortem brain tissue (ROSMAP, Religious Orders Study/Memory and Aging Project) and used BHLHE40-/41-microglia to identify microglia in the ARM state^14^. Separating the data by APOE haplotype, we found that APOE44 individuals had significantly fewer ARMs compared to APOE23 control, APOE33 control and APOE33 AD individuals (Figure 2D). Intriguingly, we saw that APOE23 AD individuals also had a decrease in ARMs and that APOE23 control individuals had an increased number of ARMs, indicating that this state change may in fact be beneficial (Figure 2D). The discrepancy between these findings and our APOE23 AD iMG results likely reflects the cell-autonomous nature of our model, which does not capture the influence of the broader cellular and systemic environment present *in vivo*. Because ARMs have been implicated in lipid clearance^14^, we asked if metabolic pathways were impaired in APOE44 iMG. To investigate this, we performed Compass^41^, an *in silico* approach to infer metabolic flux of cells based on single-cell RNA sequencing. We compared APOE44 to APOE33 iMG and observed negative Cohen’s D for the majority of predicted reactions in APOE44 relative to APOE33, indicating systemic differences in metabolic utilization between the two, including FAO, cholesterol metabolism and citric acid cycle (Figures 2E, S2C).

**Figure 2.**
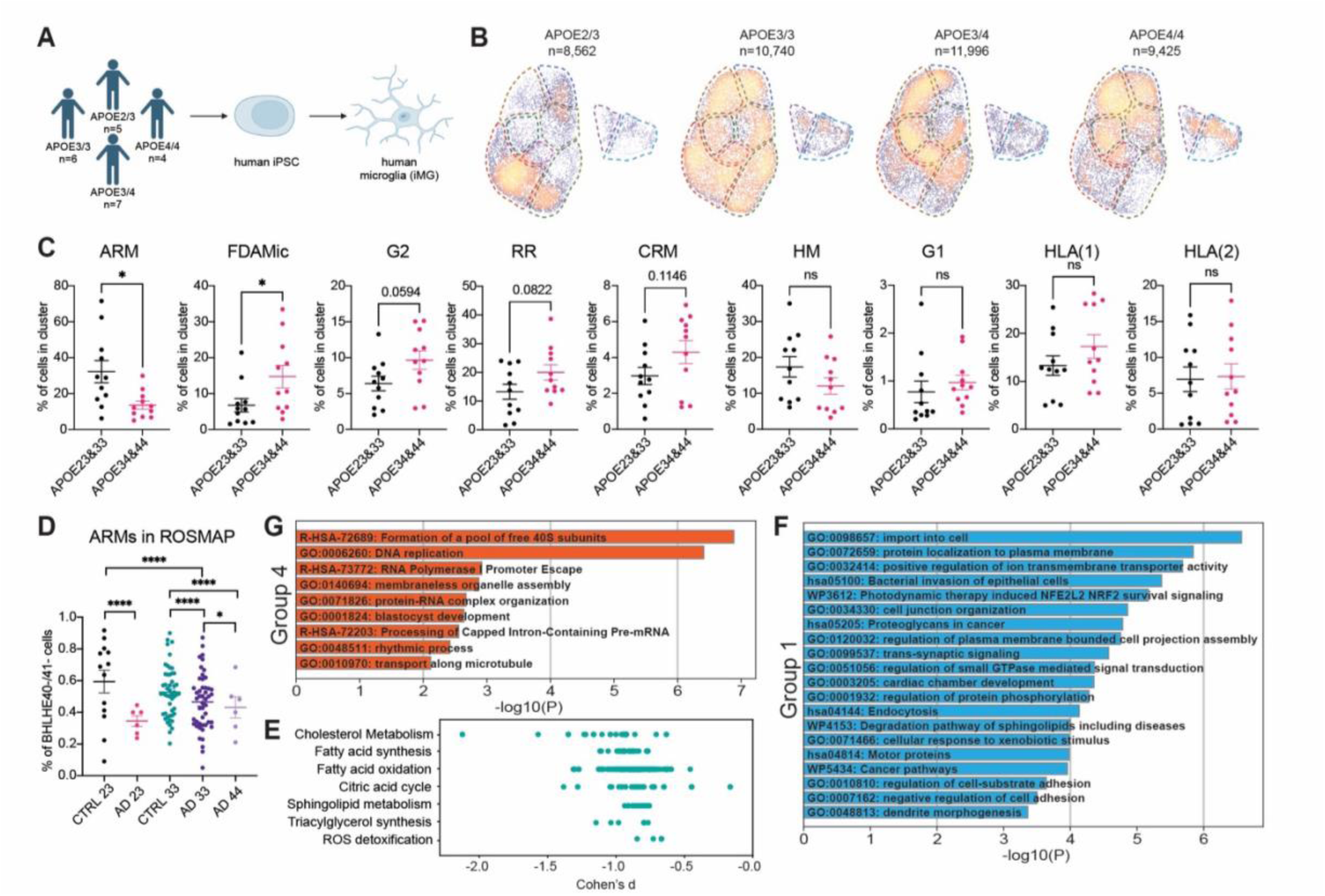
APOE4 influences state shifts in microglia, suppressing the ARM signature. (A) Schematic of donors divided by APOE haplotypes (APOE23 n = 5, APOE33 n = 6, APOE34 n = 7, APOE44 n = 4). (B) Density plot of scRNAseq UMAP per APOE haplotype displaying distribution across identified clusters. (C) Distribution and proportion of cells across different cell states, separated by APOE4-and APOE4+. Each dot represents an individual line. Unpaired t-test, two-tailed, mean ± SEM. (D) Percentage of BHLHE40-/41- microglia in ROSMAP data set, split by APOE haplotypes and diagnoses. Each dot represents an individual donor (CTRL APOE23 n = 13, AD APOE23 n = 6, CTRL APOE33 n = 50, AD APOE33 n = 48, AD APOE44 n = 6). Chi-squared test, mean ± SEM. (E) Compass analysis of metabolic pathways in APOE44 iMG vs APOE33 iMG in G0 phase cells. Select terms shown. Wilcoxon signed-rank test, significance set to padj < 0.05. (F) Enrichment analysis of downregulated genes in APOE44 compared to other haplotypes (Group 1, N=186 genes). (G) Enrichment analysis of upregulated genes in APOE44 compared to other haplotypes (Group 4, N=80 genes). Created using Metascape^97^. (C-D) *p < 0.05; ns, non-significant See also Figure S2.

Differential expression analysis across APOE groups in iMG revealed four groups of genes with distinct profiles of expression in APOE44 samples. Group 1 includes genes (N=186) that are highly expressed in APOE23 iMG and low in APOE44 iMG, creating a linear downward trend (Figure S2D). Pathway analysis revealed terms associated with reduced expression in endosomal and lysosomal activity (i.e., GM2A, SCARB2) (Figures 2F, S2E, Table S3). Group 4 is enriched for translation-related genes (N=80) that are upregulated in APOE44 (Figures 2G, S2D-E, Table S3). Sex-stratified analysis revealed marked differences in the transcriptional response to APOE haplotype. Females exhibited a substantially greater number of differentially expressed genes (n=1,513) compared to males (n=140), with only 44 genes overlapping between sexes (Figure S2F). These findings suggest a heightened transcriptional sensitivity to APOE4 in females. Gene enrichment analysis further indicated that females drive the majority of pathway-level changes observed in the combined analysis (Figures S2G-H).

We subsampled 8 APOE33 individuals and 8 APOE44 individuals from the ROSMAP study and performed pseudobulk differential analysis to compare against our iMG dataset. Similar to our findings in the iMG model, the dysregulated genes in postmortem brain tissue of APOE44 samples revealed alterations in ribosomal, lysosomal and mitochondrial pathways in microglia (Figures S2I-K, Table S4). We compared the directionality of genes identified in Groups 1 and 4 between our scRNA-seq dataset and ROSMAP microglia and observed a modest correlation (R² = 0.144) (Figures S2L-N). The overlap in pathway-level signals supports shared biological processes, and our system enables mechanistic interrogation of how APOE4 modifies these pathways. The lysosome, ER, and mitochondria form a close physical connection at specialized contact sites, allowing for direct transfer of molecules like calcium ions, lipids, and metabolites between them^44–49^. With these findings, we sought to understand how APOE4, a lipid transporter, might impact the lysosomal-ER-mitochondrial axis.

### APOE44 microglia accumulate cholesterol esters in the lysosomes, modulating lysosomal acidification

To exclude the metabolic differences between genetically distinct individuals from the impact of the APOE44 genotype on metabolism, we converted isogenic APOE33 and APOE44 iPSCs (Jackson Laboratory, JIPSC001000, JIPSC001150) into iMG, as previously described (Figure 3A)^27^. As both our Compass and GO analyses indicated there were changes in lipid metabolism in APOE44, we began by measuring lipid composition in the iMG using high-resolution mass spectrometry coupled to liquid chromatography (LC-MS). At baseline, APOE44 iMG have significantly increased lipid species compared to APOE33 iMG, with cholesterol esters (ChEs) being the most significant species (Figures 3B,3C, Table S5). Interestingly, we found that hexosylceramides (HexCer) and lactosylceramides (LacCer) were significantly lower in APOE44 microglia, which may be an indication of impaired autophagy (Figure 3D)^48^. ChEs are typically shuttled into cells via apolipoproteins, where they are hydrolyzed in the lysosomes and converted into free cholesterol^49^. This free cholesterol is then shuttled to the ER, where it provides lipid support to different parts of the cell, including the mitochondria via the mitochondrial associated membrane (MAM)^50,51^. Excess free cholesterol is esterified and packaged into lipid droplets via acetyl-CoA acetyltransferase 1 (ACAT1) for storage^52^.

**Figure 3.**
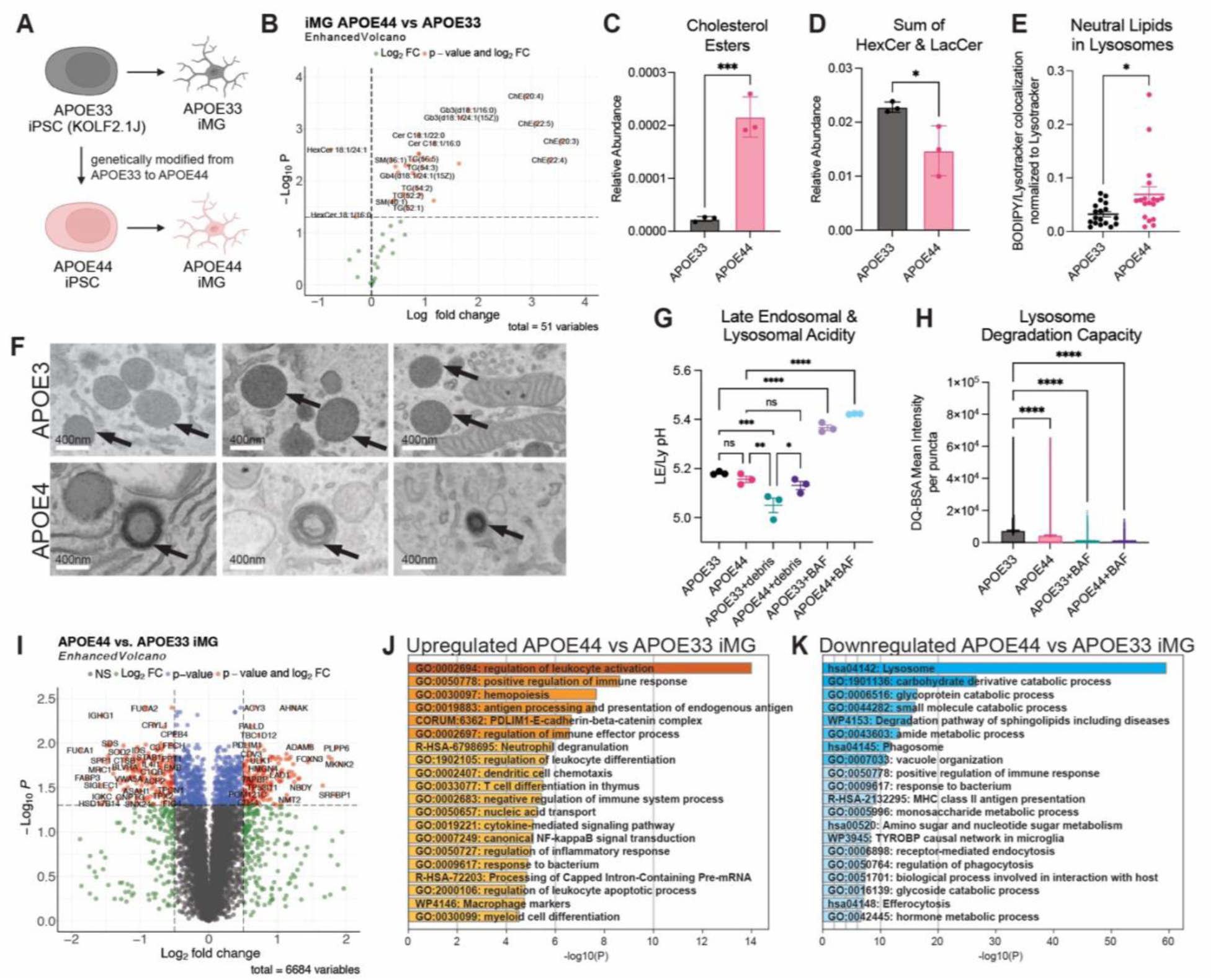
APOE44 microglia accumulate cholesterol esters in the lysosomes, modulating lysosomal acidification. (A) Schematic of APOE33 and APOE44 isogenic iMG. (B) Volcano plot of relative abundance of lipids (n=3 wells each). Significance set to p < 0.05. (C) Relative abundance of cholesterol esters (n=3 wells each). (D) Relative abundance of sum of hexosyl ceramides and lactosyl ceramides (n=3 wells each). (E) BODIPY-493/503 and LysoTracker Red DND-99 colocalization area normalized to Lysotracker area across APOE33 and APOE44 iMG. Each dot represents one frame (n = 2 wells x 9 frames per line).(F) Electron microscopy of APOE33 and APOE44 iMG focused on lysosomes. Arrows indicate representative lysosome. (G) Late endosomal and lysosomal acidity measured ratiometrically by dextran polymers labeled with the pH sensor ApHID and Alexa 647 (pH-independent) across APOE33 and APOE44 iMG untreated, treated with organoid debris or with Bafilomycin (0.1 nM) (n=3 wells each, 9 frames per well). (H) DQ-BSA to assess lysosome degradation capacity with or without Bafilomycin (10 nM). Each dot represents mean intensity per puncta (for each group, 3 wells were imaged, 9 frames each). One way ANOVA, mean ± SEM. (I) Volcano plot visualizing differential protein expression in APOE44 vs APOE33 iMG (n=3 wells each). (J,K) Enrichment analysis of upregulated (J) and downregulated (K) genes in APOE44 compared to APOE33 iMG, significance set to padj < 0.05, |Log2FC cutoff| > 0.5. Created using Metascape^97^. (B-E) Unpaired t-test, two-tailed, mean ± SEM. (G, H) One way ANOVA, mean ± SEM. (I-K) Unpaired t-test, two-tailed, mean ± SEM, corrected with Benjamini-Hochberg. *p < 0.05; **p < 0.01; ***p < 0.001; ****p < 0.0001, ns, non-significant See also Figure S3.

To visualize neutral lipids, we used the BODIPY dye and found that APOE44 iMG have significantly more puncta per cell, indicative of increased neutral lipid accumulation (Figure S3A-B). We next asked if the increase in ChEs in APOE44 iMG could also be attributed to a buildup of lipids in the lysosomes. To assess this possibility, we quantified colocalization of BODIPY and LysoTracker and found that APOE44 iMG have significantly more neutral lipid accumulation in the lysosomes (Figure 3E). This finding suggests that, in APOE44 iMG, there may be a dysfunction in the hydrolysis of ChEs in the lysosomes, resulting in accumulation of ChEs due to incomplete degradation. Another contributing factor could be that lipids are not properly secreted as loaded apolipoproteins, causing excess lipids to be shuttled to lysosomes. As an orthogonal approach to visualize lysosomes, we performed transmission electron microscopy on iMG from APOE3 and APOE4 donor lines. Indeed, we found instances in which lysosomes from APOE44 appeared more electron dense, with a concentric, whorled membranous structure, indicative of lipid buildup (Figure 3F). Interestingly, the observed lysosomal phenotype was similar to that in Tay Sachs and other lysosomal storage disorders^53^. To assess lysosomal acidification capacity, we measured late endosomal/lysosomal (LE/Ly) acidity following challenge with organoid debris. To measure LE/Ly pH, we labeled the organelles with dextran polymers tagged with pH-sensitive and pH-independent fluorescent probes (ApHID), allowing for ratiometric pH imaging^54^. The dextran polymers are composed of glucose units and cannot be degraded by lysosomal hydrolases, leading to stable accumulation in the organelles upon cellular uptake^54,55^. While baseline LE/Ly pH did not differ between APOE33 and APOE44 iMG (∼5.15), APOE33 iMG exhibited further acidification upon challenge (∼5.05), whereas APOE44 iMG failed to do so (Figure 3G). Consistently, lysosomal degradative capacity, assessed by DQ-BSA, was reduced in APOE44 iMG (Figure 3H)^56,57^. Together, these findings suggest that APOE33 iMG lysosomes retain the ability to adapt to challenge, whereas APOE44 iMG exhibit impaired lysosomal responsiveness.

To gain further clarity on the impact of APOE haplotype on different organelles, we performed proteomics on the isogenic APOE33 and APOE44 iMG. These data were in agreement with our scRNAseq results, also indicating that lysosomal and endosomal pathways were being downregulated (i.e., LAMP1, LAMP2, HEXA, HEXB) (Figures 3I-K, S3C, Table S6). Of the 84 identified lysosomal hydrolases and V-ATPases, 53 were significantly downregulated in APOE44 iMG, supporting lysosomal impairment (Figure S3D). Further, p62 (SQSTM1) and LC3B (MAP1LC3B2) were both upregulated, supporting a buildup of autophagosomes (Figure S3E). Proteins related to protein synthesis and trafficking were also dysregulated, suggesting ER stress (RPS24, RPS17, SEC24D) (Figure S3F)^58–60^. We observed that proteins important in MAM function, such as VDAC2 and HSPA9 (GRP75) were downregulated in APOE44 iMG implying a dysregulation of exchange between the ER and the mitochondria (Figure S3F)^44,45,61–63^. Taken together, these results suggest deficits in vesicle trafficking, protein homeostasis, ER function and mitochondrial-ER association. To understand if the ER-mitochondria axis is being impacted by this impairment, we were interested to see if the bioenergetics of APOE44 microglia was affected.

### APOE44 microglia shift away from FAO towards glycolysis and are proinflammatory

To understand the bioenergetics of APOE44 microglia, we first performed the Mito Stress Test Kit on the Seahorse XFe96 Analyzer using the isogenic APOE iMG (Figure 4A). We observed that APOE44 iMG had decreases in oxygen consumption rate at baseline, ATP-linked respiration and max respiration compared to APOE33 iMG (Figures 4A,4B). This result might suggest dysfunctional mitochondria and therefore we tested their function. We did not observe differences in proton leak, spare capacity, or non-mitochondrial oxygen consumption, although there was a downward trend (Figure S4A). Because we also observed an increase in LDHA (lactate dehydrogenase) gene expression levels in APOE44 donor iMG (Figure S4B), we were interested in whether APOE44 microglia shifted their bioenergetics from OXPHOS to glycolysis. We quantified the relative abundance of lactate in iMG and found that APOE44 microglia had significantly larger intracellular pools (Figures 4C, Figure S4C). We measured secreted lactate in the media as well and observed an increase in lactate efflux from the APOE44 microglia (Figure 4D). Treating cells with [U-^13^C_6_] glucose, we quantified incorporation into the TCA cycle by GC-MS in APOE33 and APOE44 iMG. We found that there was no difference in the incorporation of glucose into citrate and other TCA metabolites after 24 hrs (Figure 4E, S4D). Microglia are known to utilize FAO, a process of breaking down fatty acids for energy, so we traced the incorporation of [U-^13^C_16_] palmitate into the TCA cycle^34^. We observed a decrease in the mole percent enrichment (MPE) of citrate, malate, glutamate and aspartate in APOE44 iMG compared to APOE33 iMG (Figure 4F). This finding suggests a decreased ability to catabolize lipids.

**Figure 4.**
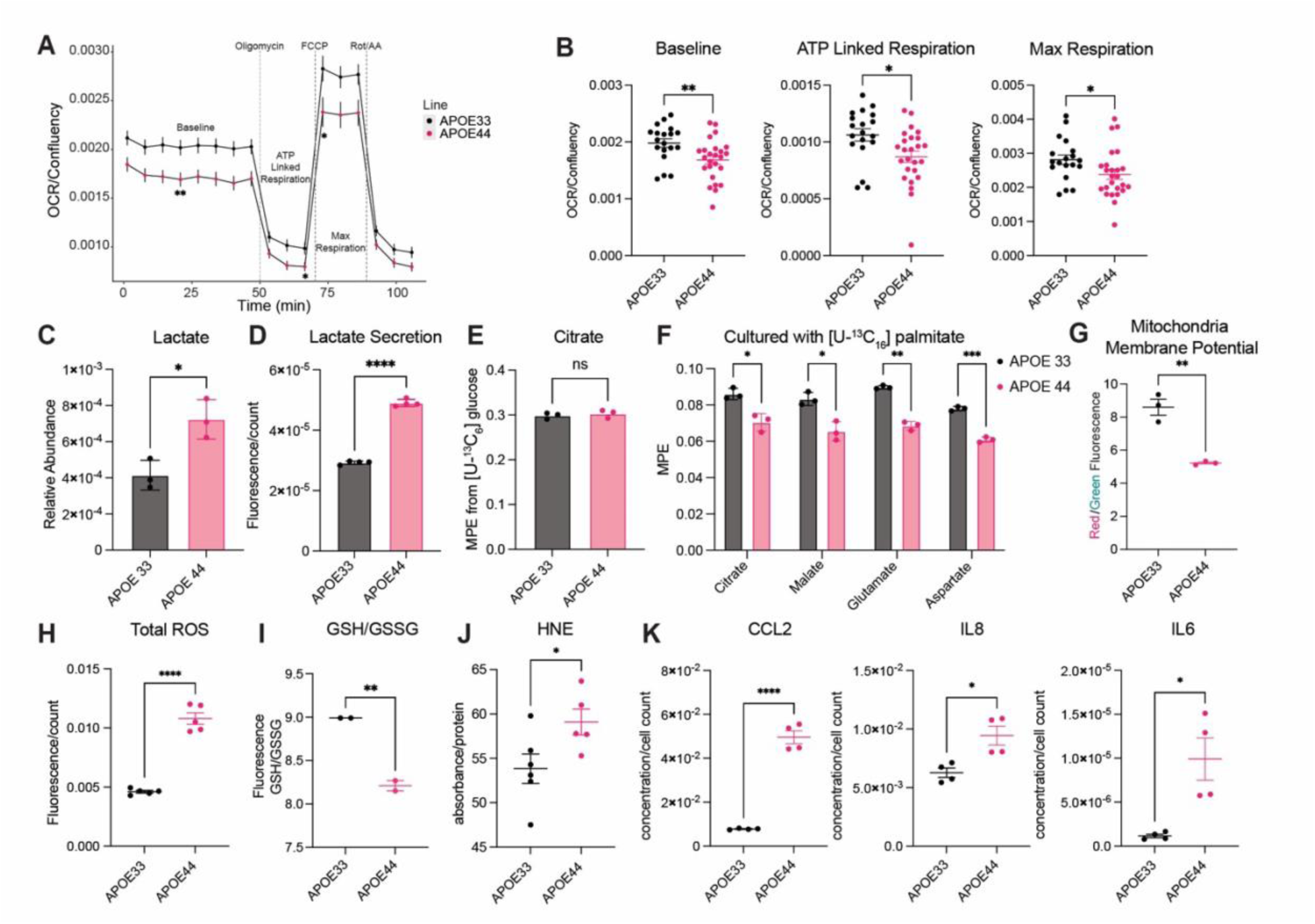
APOE44 microglia shift away from FAO towards glycolysis and are proinflammatory. (A) Seahorse XFe96 Mito Stress Test Kit of APOE33 and APOE44 iMG with different drug treatments shown. APOE33 iMG n = 19 wells, APOE44 iMG n = 25 wells. (B) Baseline oxygen consumption rate, ATP linked respiration, and max respiration normalized to confluency of wells, comparing APOE33 and APOE44 iMG. Each dot represents a replicate well. (C) Relative abundance of lactate after tracing with [U-^13^C_16_] palmitate. (D) Lactate secretion measured in supernatant using kit (ab65330), normalizing to cell count. (E) Labeled citrate after [U-^13^C_6_] glucose tracing. (F) Labeled citrate, malate, glutamate, and aspartate after [U-^13^C_16_] palmitate tracing. (G) Mitochondria membrane potential (MMP) measured with JC-1 dye comparing APOE33 and APOE44 iMG. On a Glomax plate reader, red was measured at an excitation of 520 nm and green at an excitation of 475 nm. (H) Measurement of superoxides and hydroxyl radicals, normalized to cell count. (I) GSH/GSSG levels of cell lysates measured by kit (MedChem, HY-K0311). (J) Measurement of HNE across APOE33 and APOE44 iMG normalized to protein. (K) Cytokine analysis of CCL2, IL8 and IL6 from media collected from APOE33 and APOE44 iMG, normalized to cell count. (A-D, G-K) Unpaired t-test, two-tailed, mean ± SEM. (E-F) Two-way ANOVA, Šídák’s multiple comparisons test, mean ± SEM. *p < 0.05; **p < 0.01; ***p < 0.001; ****p < 0.0001 See also Figure S4.

To further assess the health of the mitochondria, we measured mitochondria membrane potential (MMP) and observed a significant reduction in APOE44 iMG, typically a sign of dysfunctional mitochondria (Figures 4G, S4E)^64^. Given this result, we next measured ROS and found that APOE44 iMG produced significantly more ROS, specifically superoxides and hydroxyl radicals, compared to APOE33 iMG, which could be a response to low MMP (Figure 4H)^65^. APOE44 iMG exhibited a decreased GSH/GSSG ratio, but no change in total glutathione, indicative of reduced ROS scavenging (Figure 4I, S4F). This finding is supported by our proteomics data revealing a reduction of proteins important in detoxification like SOD2, NQO1 and SQOR (Figure S4G).

Given the role of APOE in lipid regulation, we asked if increased ROS resulted in increased lipid oxidation. Importantly, oxidized low density lipoproteins (oxLDL) are less likely to be hydrolyzed in the lysosome^66,67^. Our scRNAseq showed that ALOX5AP (arachidonate 5-lipoxygenase activating protein), a microglia-specific dioxygenase that catalyzes the peroxidation of polyunsaturated fatty acids and promotes inflammation, was significantly upregulated in APOE44 donor iMG (Figure S4H)^68,69^. To detect lipid oxidation, we measured HNE (4-hydroxy 2-nonenal), a byproduct of lipid oxidation, in the isogenic APOE33 and APOE44 iMG^70^. APOE44 iMG had significantly increased HNE compared to APOE33 microglia (Figure 4J). These results agree with our *in vitro* RNAseq data and with the *in vivo* postmortem data (ROSMAP), where AD individuals had an increase in the percentage of microglia expressing ALOX5AP compared to control individuals (Figure S4I). Microglia that have shifted towards glycolysis and secrete ROS are associated with a proinflammatory state in microglia^71,72^; therefore, we measured key cytokines in our isogenic iMG and found that CCL2, IL-6 and IL-8 were all increased in APOE44 iMG (Figure 4K). Taken together, these findings indicate APOE44 iMG have reduced capacity to utilize FAO and increased glycolytic flux due to dysfunctional mitochondria, which is accompanied by a proinflammatory state.

### APOE4 microglia exhibit an increased population of G2 senescent-like cells

Dysfunctional mitochondria, proinflammatory cytokine release and lipid accumulation are typically features of senescence; therefore, we aimed to understand whether APOE44 contributed to microglia shifting to a senescent-like state^73^. While comparing various microglia datasets with ours, we discovered the lipid-high microglia signature from Haney et al. overlapped with our G2 cluster (Figure 5A)^10^. This cluster expresses known senescent markers CDKN2A (p16) and CDKN1A (p21) (Figure 5B), and G2 cell cycle markers like KI67, TOP2A, and CENPF, and DNA damage/replication genes like RRM1, BUB1B, and H2AFX (Figure S5A,S5B,S5C)^74,75^. We performed cell cycle prediction to identify the phase of the cells more accurately and observed a decrease in APOE4 iMG in G0 and an increase in G2 (Figure 5C). CDKN2A (p16) staining revealed that p16+ iMG had increased nucleus size, a known feature of senescence (Figures 5D,5E)^76^. We sought to understand if these cells had been arrested in a G2 cell cycle phase and left in a G2 senescent-like state. G2 cell cycle arrest leaves cells in a G1 tetraploid state; therefore, we isolated this population for further analysis^77^. Using the Vybrant DyeCycle dye with verapamil to avoid dye efflux, we isolated cells with 4N DNA content from APOE44 microglia using FACS (Figures 5F, S5D). Seventy-two hours after plating the isolated 4N microglia, the cells were reanalyzed for DNA content. Cells originally designated G0 remained 2N, while cells originally designated G2 remained 4N, confirming that these cells had not divided and were arrested in a tetraploid state (Figure 5G). Based on our observation from our gene module analysis, we tested if lipid content was higher in this G2 senescent-like population. Lipidomics revealed that G2-arrested microglia had significantly increased ChEs relative to G0 microglia (Figure 5H). Further, data from postmortem brain tissue (ROSMAP) also demonstrated that AD individuals had increased p16+ microglia and that APOE44 AD individuals had significantly more than APOE33 AD individuals, highlighting APOE44’s role in senescence (Figure 5I). To analyze these results based on cell cycle phase, we looked at p16+ microglia separated by predefined ROSMAP clusters and found that the MKI67+ cluster, or the G2 cluster, contained a larger percentage of the p16+ microglia compared to the other two clusters, supporting our finding that G2 senescent-like microglia can occur *in vivo* (Figure S5E).

**Figure 5.**
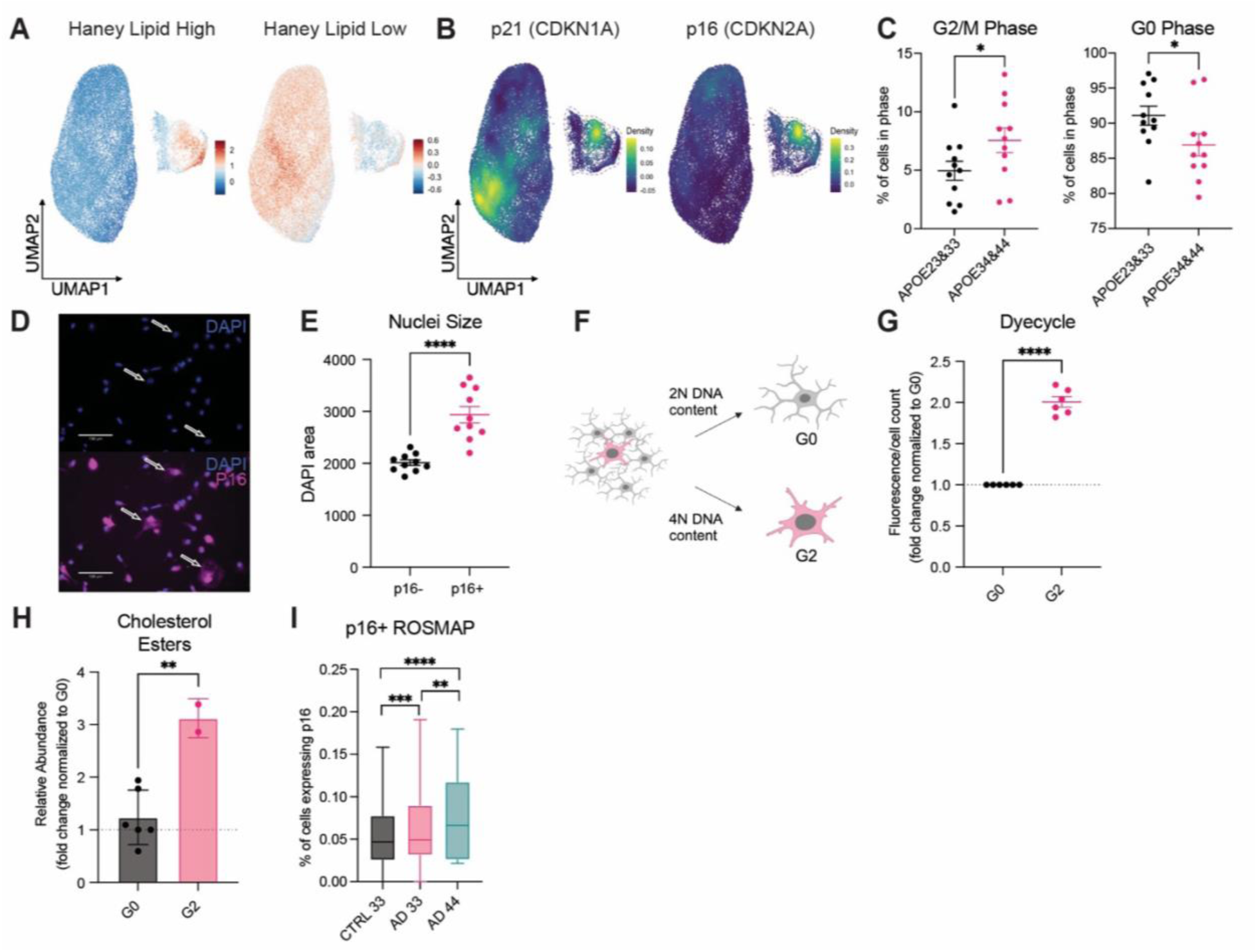
APOE4 microglia exhibit an increased population of G2 arrested cells. (A) Module scores created using DEGs from Haney et al. plotted on our UMAP space^10^. Left represents upregulated genes in lipid high APOE44 iMG, right represents downregulated genes in lipid high APOE44 iMG. (B) Density UMAP of expression of CDKN1A (p21) and CDKN2A (p16). (C) Distribution and proportion of cells in G2/M and G0 phase, separated by APOE4- and APOE4+. Each dot represents an individual line. Mann-Whitney U-test, two-tailed, mean ± SEM. (D) Representative image of p16 staining with DAPI on iMG. Scale bar = 130µm. Arrows point toward examples of nuclei with larger areas. (E) ImageJ analysis of nuclei size measuring area of DAPI from p16- and p16+ iMG. Each dot represents a different cell (p16- n = 10, p16+ n = 10). (F) Schematic of using Vybrant DyeCycle Violet Dye with verapamil to stain live nuclei in APOE44 iMG to collect 2N (G0) and 4N (G2) cells. (G) Using a plate reader, G0 and G2 cells that were collected were stained again with DyeCycle and verapamil 72 hours later. G2 is normalized to G0 in each run and cell count. Each dot represents a replicate well. (H) Relative abundance of cholesterol esters between G0 and G2 cells normalized to G0. (I) p16+ microglia in ROSMAP dataset across different APOE haplotypes and diagnoses. CTRL APOE33 n = 50, AD APOE33 n = 48, AD APOE44 n = 6. (E, G, H) Unpaired t-test, two-tailed, mean ± SEM. (I) Chi-squared test, mean ± SEM. *p < 0.05; **p < 0.01; ***p < 0.001; ****p < 0.0001. ns, non-significant See also Figure S5.

### LXR agonist GW3965 is insufficient to alleviate APOE44 iMG dysfunction

LXR agonists have been used as rescue molecules in several APOE4 studies; therefore, we were interested in how this drug would impact lipid metabolism and bioenergetics^7,8^. We treated our APOE44 iMG with the GW3965 LXR agonist for 24 hours at 10 µM (Figure 6A) and found that the MMP in APOE44 iMG treated with GW3965 was significantly increased (Figure 6B). Next, we measured cytokine secretion and found that CCL2 and IL6 levels were decreased with treatment (Figure 6C), although IL8 was not rescued and instead increased. We performed bulk RNAseq (Figure 6D), and PCA analyses revealed that the largest source of variability in the dataset was treatment. Interestingly, APOE44 iMG treated with GW3965 and APOE33 iMG moved closer together on the secondary axis of variation, defined by G1 phase genes and immune response genes (Figure S6A, Table S7). GW3965 treatment reduced inflammatory gene markers and increased lipid metabolism genes (Figures 6E, 6F, Table S7). Unbiased hierarchical clustering analysis demonstrated that APOE44 iMG treated with GW3965 were relatively equidistant from APOE33 iMG and APOE44 iMG treated with DMSO, supporting a shift of APOE44 back towards APOE33 (Figure S6B), although significant differences remain between the two, indicating that GW3965 partially rescues the transcriptomic phenotype (Figure S6C-D). We observed that NFKB1 and NFKB2, both elevated in APOE44 iMGs, are suppressed by GW3965 (Figure S6E). Mechanistically, GW3965 induces LXR SUMOylation, promoting its interaction with NFKB and thereby repressing NFKB-driven inflammatory gene expression^78^. Module scores of differentially upregulated genes overlapped the most with the ARM cluster whereas downregulated genes overlapped the most with the CRM cluster (Figure S6F). In line with prior studies suggesting that LXR-mediated upregulation of cholesterol efflux is beneficial^8^, GW3965 increased expression of the cholesterol transporters ABCA1 and ABCG1. However, this increase was accompanied by induction of lipogenic and lipid storage programs, including SREBF1 and PLIN2 (Figure S6G; Table S7). Consistent with this finding, lipidomic profiling revealed a reduction in cholesterol esters alongside an increase in triglycerides (Figures 6G).

**Figure 6.**
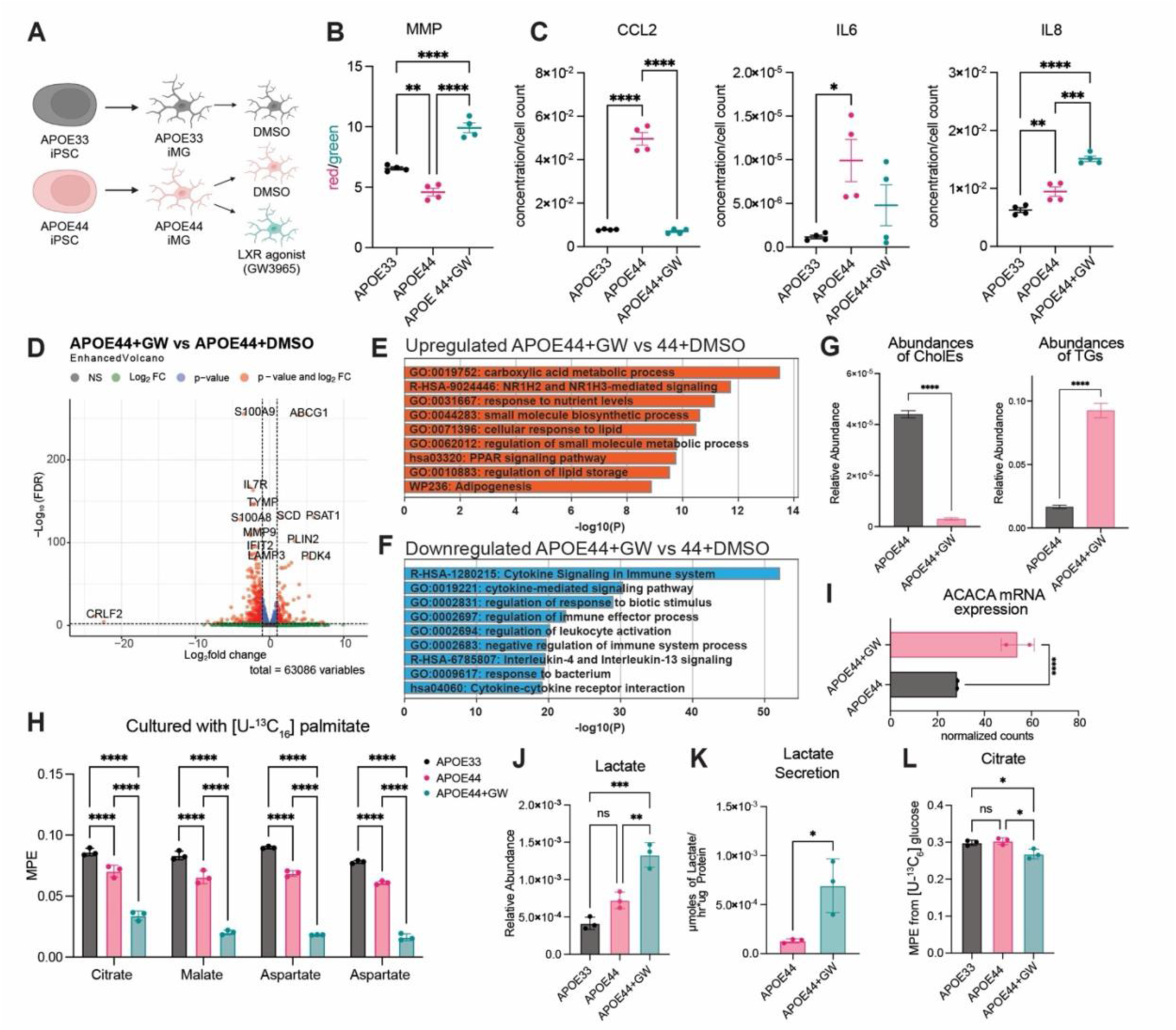
LXR agonist GW3965 is insufficient to alleviate APOE44 iMG dysfunction. (A) Schematic of APOE33 and APOE44 isogenic microglia, with APOE44 treated with LXR agonist GW3965 and controls treated with DMSO. (B) Mitochondria membrane potential (MMP) measured with JC-1 dye comparing APOE33 and APOE44 iMG. On a Glomax plate reader, red was measured at an excitation of 520 nm and green at an excitation of 475 nm. (C) Cytokine analysis of CCL2, IL6, and IL8 from supernatant, normalized to cell count. (D) Volcano plot of DEGs comparing APOE44(GW) to APOE44(DMSO). (E, F) Enrichment analyses of (E) upregulated and (F) downregulated genes in APOE44(GW) vs APOE44(DMSO). Top 9 terms are shown. Created using Metascape^97^. (G) Relative abundance of cholesterol esters and triglycerides between APOE44 iMG +DMSO and +GW3965 (n=3 wells each). (H) Mole percent enrichment of citrate, malate, glutamate, and aspartate after culture for 24 hours with [U^-13^C_16_] palmitate. (I) ACACA mRNA expression. (J) Relative abundance of lactate after culture with 100µM BSA conjugated [U-^13^C_16_] palmitate. (K) Lactate secretion measured by YSI. (L) Mole percent enrichment of citrate after culture for 24 hours with [U-^13^C_6_] glucose. (B-C) One way ANOVA, mean ± SEM. (D-F) Wald test, corrected using the Benjamini-Hochberg correction for multiple testing, significance set to padj < 0.01 and |Log2FC cutoff| > 1. (G, K) Unpaired t-test, two-tailed, mean ± SEM. (H, J, L) One way ANOVA, Tukey’s multiple comparisons test, mean ± SEM. (I) Wald test, corrected using the Benjamini-Hochberg correction for multiple testing, mean ± SEM.*p < 0.05; **p < 0.01; ***p < 0.001; ****p < 0.0001. ns, non-significant See also Figure S6.

To determine whether this shift in lipid metabolism restored FAO, we performed metabolic tracing using [U-^13^C_16_] palmitate. Unexpectedly, GW3965 treatment further reduced FAO (Figure 6H). Transcriptomic analysis revealed upregulation of ACACA (Figure 6I), which encodes acetyl-CoA carboxylase (ACCα), the rate-limiting enzyme in fatty acid synthesis. Elevated ACCα activity increases malonyl-CoA levels, which inhibit carnitine palmitoyltransferase 1 (CPT1), thereby restricting mitochondrial fatty acid import and oxidation^79^. Together, these findings indicate that LXR agonism promotes fatty acid synthesis and storage – reflected by increased SREBF1, PLIN2, and ACACA – at the expense of fatty acid utilization.

We next examined the impact of LXR activation on cellular bioenergetics. In APOE44 iMGs, GW3965 treatment increased lactate production and secretion (Figures 6J, 6K) and tracing with [U-^13^C_6_] glucose revealed a reduced incorporation of glucose into citrate (Figure 6L). Collectively, these data demonstrate that LXR agonism further drives a metabolic shift away from oxidative phosphorylation toward glycolysis, indicative of a less efficient mode of energy production.

### APOE44 microglia have impaired lipoprotein secretion, providing less lipid support to neurons

To investigate how the dysfunction we have identified in APOE44 microglia might play a role in increased risk for AD, we performed coculture experiments with fibroblast-derived iNs, which maintain the epigenetic characteristics of the original donor^80^. iMG and iNs were cocultured in a transwell system in which they were separated by a porous membrane. All iNs were derived from a healthy, aged individual with an APOE33 haplotype. These iNs were cultured with APOE33 or APOE44 isogenic iMG (Figure 7A, Table S5). Lipidomic profiling showed that coculture with iNs led to a broad increase in lipid species in APOE33 iMGs, notably cholesterol (Figures 7B, 7C, Table S5). Strikingly, the opposite pattern was observed in APOE44 iMGs, where coculture resulted in a reduction of multiple lipid species, including a significant decrease in triglycerides (Figures 7B, S7A). Abundance of ceramide C18:1/18:0 was increased in both the APOE33 and 44 iMG after coculturing with iNs, indicating this ceramide might be taken up by the iMG from the iNs (Figures 7B, S7A, Table S5). iNs cocultured with APOE33 iMG had a significant increase in numerous lipids, including sphingomyelins (SMs) and free cholesterol, when compared to iNs cultured with APOE44 iMG (Figures 7D, 7E, Table S5). Although TGs were not significantly increased in iNs cocultured with APOE33 iMG, there was an increasing trend in individual TG species and the sum of TGs (Figures 7D, 7E). These differences in lipid species in iNs after coculturing with APOE33 or APOE44 iMG suggests that APOE44 iMG are unable to provide the same level of lipid support that APOE33 iMG do. These results identify a previously unrecognized role for healthy microglia in lipoprotein production and lipid delivery to aged neurons, building on a prior developmental study of microglial lipid support^81^.

**Figure 7.**
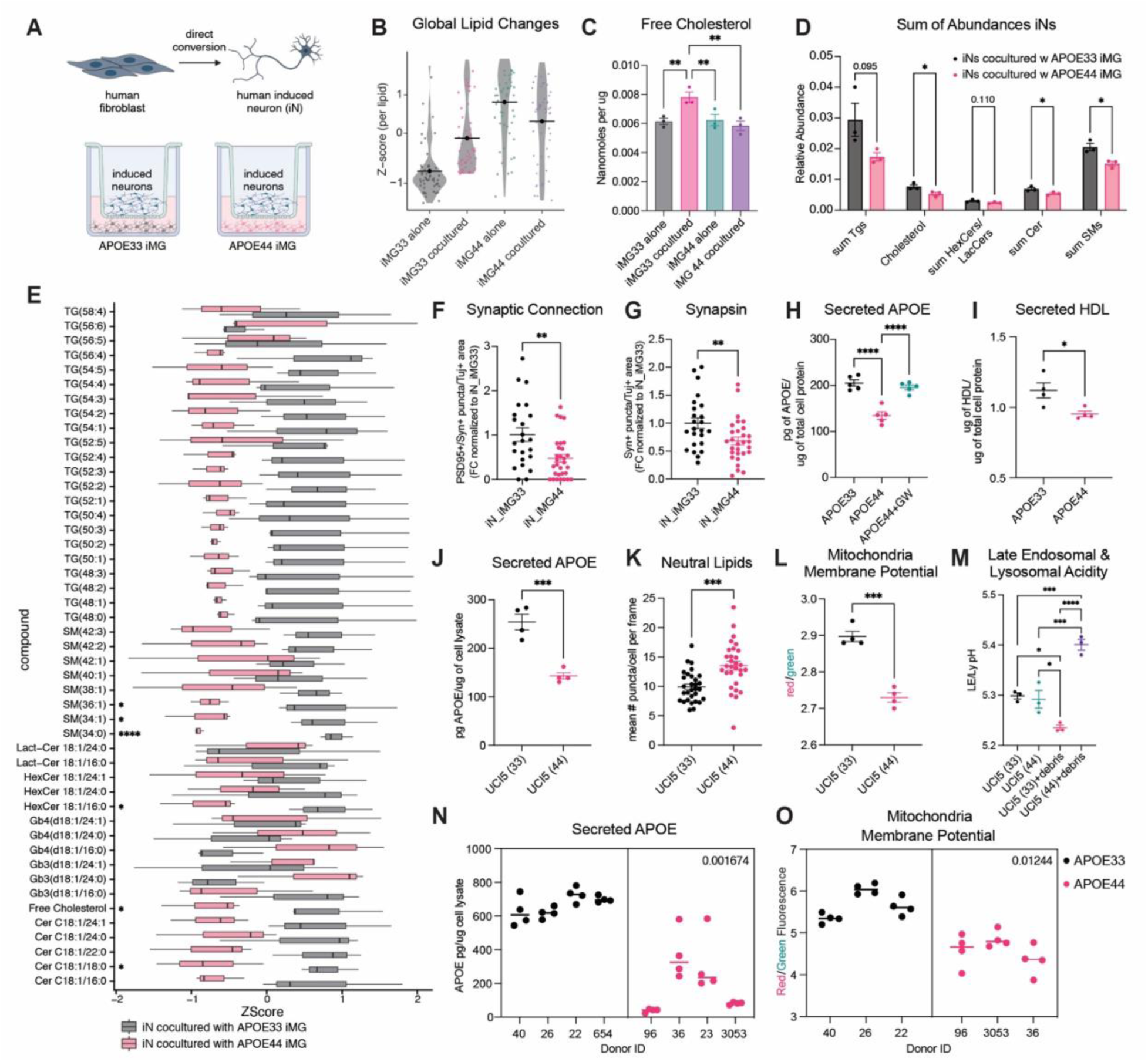
APOE44 microglia have impaired lipoprotein secretion, providing less lipid support to neurons. A) Schematic of transwell setup of induced neurons (iNs) with APOE33 and APOE44 iMG. (B) Violin plot of lipidomics of APOE33 and APOE44 iMG cocultured with iNs or alone. Each dot represents a different lipid species within the iMG lysates. (C) Free cholesterol compared between APOE33 and 44 iMG, cocultured with iNs or alone. (D) Sum of lipid species in different lipid classes compared between iNs cocultured with APOE33 iMG vs with APOE44 iMG. (E) Boxplot of ZScores of lipid species from iNs cocultured with APOE33 iMG and APOE44 iMG. (F) PSD95+/Synapsin+ puncta and (G) Syn+ puncta quantified in iNs cocultured with either APOE33 or APOE44 iMG, normalized to area of Tuj1. Plotted as fold change differences normalized to iN cocultured with APOE33 iMG. (H) APOE ELISA of supernatant from APOE33 and APOE44 iMG (n = 5 wells each). (I) HDL-like particles measured in supernatant and normalized to protein of lysate (n = 4 wells each). (J) APOE ELISA of supernatant from UCI5 APOE33 and UCI5 APOE44 iMG (n = 4 wells each). (K) BODIPY-493/503 quantified as mean number of puncta per cell per frame across UCI5 APOE33 and APOE44 iMG (n = 3 wells x 10 frames per group). (L) MMP measured with JC-1 dye comparing UCI5 APOE33 and UCI5 APOE44 iMG (n = 4 wells each). On a Glomax plate reader, red was measured at an excitation of 520 nm and green at an excitation of 475 nm. (M) Late endosomal and lysosomal acidity measured ratiometrically by dextran polymers labeled with the pH sensor ApHID and Alexa 647 (pH-independent) across UCI5 APOE33 and APOE44 iMG untreated and treated with organoid debris (n=3 wells each, 25 frames per well). (N) APOE ELISA of supernatant from 8 donor lines, 4 APOE33 lines and 4 APOE44 lines (n = 4 wells each). (O) MMP measured with JC-1 dye comparing 6 donor lines, 3 APOE33 lines and 3 APOE44 lines (n = 4 wells each). On a Glomax plate reader, red was measured at an excitation of 520 nm and green at an excitation of 475 nm. (D-G, I-M) Unpaired t-test, two-tailed, mean ± SEM. (C, H) One way ANOVA, mean ± SEM. (N, O) Nested t-test, two-tailed, mean ± SEM. *p < 0.05; **p < 0.01; ***p < 0.001; ****p < 0.0001 See also Figure S7.

To evaluate the functional consequences of reduced lipid support, we co-cultured iNs with APOE33 or APOE44 iMG and stained for PSD95, Synapsin, and Tuj1 (Figure S7B). iNs co-cultured with APOE44 iMG displayed reduced Synapsin puncta density and decreased PSD95/Synapsin colocalization (normalized to Tuj1 area), indicating impaired synaptic formation (Figure 7F, 7G). These findings align with established roles of lipids in supporting synaptic function and maintenance^82–84^. To understand if APOE44 iMG have reduced apolipoprotein secretion, we measured APOE protein in lysates and supernatant of APOE33 and APOE44 iMG. We found that APOE44 iMG produced and secreted significantly less APOE in comparison to APOE33 iMG, which we were able to rescue with the LXR agonist GW3965; this rescue could be due to the increase in APOE mRNA with GW3965 (Figures 7H, S7C-D). We then measured HDL-like particles in the supernatant and found significantly fewer particles in APOE44 iMG (Figures 7I). Together, these data suggest that APOE44 iMGs have a reduced capacity to secrete lipoproteins compared to APOE33 iMGs, thereby limiting their ability to provide lipid support to neurons.

To assess the robustness of these findings, we incorporated an additional female APOE isogenic pair (UCI5) and observed consistent effects across multiple readouts, including APOE secretion, MMP, and lipid accumulation (Figures 7J-L). Upon challenge with organoid debris, UCI5 APOE44 iMG lysosomes failed to mount an adaptive acidification response and instead shifted toward alkalinization (Figure 7M). Interestingly, this line accumulated higher levels of intracellular APOE protein, consistent with impaired secretion rather than reduced synthesis (Figure S7E). Furthermore, analyses in independent donor-derived lines confirmed these results, with consistent differences observed in APOE secretion (n=4 APOE33, n=4 APOE44) and MMP (n=3 APOE33, n=3 APOE44) (Figures 7N, 7O).

We performed bulk RNA-seq and proteomic profiling on the additional APOE isogenic pair (UCI5) and compared these data to our previously characterized KOLF line, which is male. Upregulated proteins were related to MHC Class II response and inflammation, whereas downregulated proteins were related to DNA replication (Figure S7F-G). Notably, we observed minimal concordance in the directionality of transcripts and proteins across the two pairs, as reflected by a low coefficient of determination (Figure S7H-I). While these differences may be influenced by sex, the limited number of lines precludes definitive conclusions. Bulk RNA-seq also lacks the resolution to distinguish microglial states, underscoring the value of single-cell approaches. Nevertheless, both lines at the proteomic level show an increase in MHC Class II response and inflammatory response and a decrease in lysosomal proteins (Figure S7I). Additionally, directionality of transcripts and proteins between APOE44 and APOE33 iMGs within each pair showed limited overlap (Figures S7J-K), highlighting the importance of functional assays to contextualize molecular changes. Within each pair, transcript-protein concordance was moderate, with 53.8% (3,589/6,676) agreement in KOLF and 60.9% (4,423/7,262) in UCI5. Given prior reports of generally low correlation between transcriptomic and proteomic datasets^85^, these observations are not unexpected. These findings motivate deeper exploration into APOE4-dependent sex-specific mechanisms.

## DISCUSSION

We provide a transcriptomic dataset of iPSC-derived microglia with different APOE haplotypes, allowing us to investigate the impact that APOE has on state shifts. APOE44 carriers have an increased risk of developing late onset AD, and a recent study highlights the pathological differences in APOE44 carriers, with many showing amyloid beta plaques^86^. The APOE protein is highly expressed in glia and is a key transporter of lipids, specifically in the form of lipoproteins^87^. Lipoproteins contain mainly cholesterol esters and triglycerides in the core but also contain free cholesterol and other phospholipids on the surface, like SM and phosphatidylcholine^88^. By studying microglia *in vitro* without perturbations, we begin uncovering the cell autonomous roles APOE4 plays in microglia. We demonstrate that APOE4 suppresses the ARM state, a population previously thought to be absent in unperturbed *in vitro* systems^29^. Importantly, this state is distinct from pro-inflammatory microglial states, underscoring the need to clearly differentiate between these functionally divergent populations. Existing human postmortem datasets support our finding that APOE4 suppress a shift toward ARMs, even in the presence of AD pathology^89,90^, supporting our idea that ARMs are neuroprotective. Our scRNA-seq dataset and the ROSMAP scRNA-seq dataset showed a modest correlation; this level of concordance is expected given fundamental differences between the systems. Our model isolates cell-autonomous effects in a controlled human context, whereas postmortem datasets reflect complex influences from other brain cell types and peripheral factors such as age, injury, infection, and diet. Additionally, ROSMAP is based on single-nucleus RNA-seq, whereas our data are derived from whole cells; prior studies have shown that single-nucleus approaches can underrepresent key microglial activation genes, including APOE, SPP1, and CD74^91^. Despite these differences, the overlap in pathway-level signals supports shared biological processes which encouraged further investigation.

We found that neutral lipids accumulate in APOE44 iMG, not only within lipid droplets, but also in lysosomes. This accumulation drives lysosomal dysfunction, resulting in impaired degradative capacity and a reduced ability to dynamically acidify in response to stress relative to APOE33.. Other studies have identified triglycerides as the main accumulating lipid in APOE44 microglia, whereas our study points to ChEs^10,92^. While lipid dyes are helpful in identifying neutral lipids, they do not differentiate between lipids such as TGs and ChEs, therefore, lipidomics should be performed. Additionally, identifying the location of these lipids, whether in droplets or accumulated in organelles such as lysosomes, is crucial to understanding the extent of the dysfunction.

Studies have shown that microglia proliferate in response to injury and pathology, which can lead to senescence^25,26^. In our study, we found that APOE4 promoted a population of G2 cell cycle-arrested iMG expressing senescent markers p21 and p16 and they have increased cholesterol ester accumulation compared to G0 cells. G2 senescence has previously been overlooked, and our study highlights the importance of creating new signature sets for different types of senescence.

Studies have shown that cholesterol is crucial at the MAM, which tethers the ER and mitochondria^46,93^. Altering cholesterol, which alters the MAM distance, leads to dysfunctional mitochondria producing less ATP and a shift towards glycolysis^94^. We also observed that APOE44 iMG have reduced FAO and shift towards glycolysis. Our data support a model in which APOE44 iMGs exhibit disrupted ER-to-mitochondria lipid transfer, driven by chronic diversion of lipoprotein-derived lipids to lysosomes and their subsequent redistribution to the ER.

While GW3965 partially rescues select APOE44 iMG phenotypes, our data suggest that LXR agonism is not a suitable therapeutic strategy in this context. Although LXR activation increases cholesterol efflux pathways (e.g., ABCA1, ABCG1), it simultaneously drives lipogenesis and lipid storage programs (e.g., SREBF1, ACACA), reduces fatty acid oxidation, and shifts metabolism toward glycolysis. Consistent with liver studies showing increased VLDL production, LXR activation appears to favor lipid packaging and export rather than utilization. In APOE4 microglia, where lipid handling is already impaired, this lipid production and packaging induced by LXR activation exacerbates metabolic dysfunction. Together, these findings suggest that broad LXR activation reinforces maladaptive lipid and bioenergetic states, highlighting the need for more targeted approaches.

We propose that microglia produce and secrete lipoproteins to support lipid homeostasis in neighboring cells, including neurons. Our data show that APOE44 microglia provide diminished lipid support compared to APOE33, driven by impaired apolipoprotein secretion. This defect leads to intracellular lipid accumulation, further exacerbating microglial dysfunction. We believe that lysosomes attempt to degrade the lipoproteins that accumulate within the cells and eventually become dysfunctional due to an excess of undegraded lipids. Lipid oxidation by the increased ROS in the system may further hinder the lysosomes’ ability to degrade these lipids. We suggest the lipid droplet accumulation is a consequence of shuttling lipoproteins to the lysosomes and that the lysosomal lipid accumulation is the main detrimental factor.

Given that aging is associated with progressive lysosomal deacidification and reduced degradative capacity, this impaired adaptability may render APOE4 microglia particularly vulnerable to age-related stress, leading to cumulative dysfunction and exacerbation of disease-relevant phenotypes over time.

Microglia are known to have less acidic lysosomes in comparison to other cells in the brain^17^. We believe this is a contributing factor as to why microglia are highly impacted by APOE4, and why others have found microglia, in particular, aggregate amyloid beta with APOE in their lysosomes^95^. Additionally, the highly phagocytic nature of microglia places considerable cargo-processing demands on lysosomes, likely exacerbating stress on this organelle. We propose a more suitable therapeutic approach is a two-pronged strategy that enhances lysosomal degradation of lipoproteins while promoting downstream mitochondrial processing of excess cholesterol into exportable metabolites, such as 27-hydroxycholesterol.

In conclusion, our study helps uncover the role APOE4 plays in microglia and subsequently in AD. The results stress the importance of stratifying donor populations by APOE haplotype when studying AD. Our data support a model in which APOE4 acts as a loss-of-function variant, rather than a gain-of-function as previously proposed^96^, leading to impaired lipid handling and cellular dysfunction. These findings highlight distinct mechanisms of pathology in APOE4 carriers and underscore the need for tailored prevention strategies. Our dataset provides a large, donor-derived population of human iMG that will help the microglia field move closer to a more complete understanding of state shifts in microglia. Additionally, we have explored how APOE44 impacts lipid metabolism and bioenergetics and how these processes impact surrounding cells. We have compared our findings to a human postmortem dataset and observed similar findings, highlighting the utilization of iPSC-derived systems to achieve relevant mechanistic insights into human disease.

## RESOURCE AVAILABILITY

## Lead contact

Further information and requests for resources and reagents should be directed to and will be fulfilled by the lead contact, Dr. Fred H. Gage.

## Materials availability

This study did not generate new unique reagents.

## Data and code availability

- All data reported in this paper will be shared by the lead contact upon request.
- All raw data will be deposited at NCBI GEO and MassIVE upon paper acceptance and will be made publicly available. This paper does not report original code.
- Any additional information required to reanalyze the data reported in this paper is available from the lead contact upon request.

## ACKNOWLEDGMENTS

We are grateful to all donors participating in this study through the UCSD and UCI Alzheimer’s Disease Research Center (ADRC). Special thanks to Gage lab members Jeffrey R. Jones, Thais S. Sabedot and M.L. Gage for editorial support, Karl Wessendorf-Rodriguez and the Metallo lab for assisting on the lipidomic studies, and Snigdha Sarkar, Rowan S. Wooldridge, and John T. Melchior for assisting with proteomics, Michael S. Cuoco for demuxlet assistance, Thais Sabedot for ROSMAP analyses, Qiang Xiao and Christina K. Lim for imaging assistance, Sarah Fernandes for the human organoids, and Jean Paul Chadarevian for the generation of the UCI ADRC5 isogenic APOE iPSCs. Thank you to Santiago Solé-Domènech, Lizzie West, Jillybeth Burgado, Bilal Kerman, Christina K. Lim, Johannes Schlachetzki, Christopher Glass, Nicola Allen, Nicole Coufal and Eduardo Macagno for the helpful discussions.

We acknowledge support from NIH grant R37-AG072502 (to F.H.G.), the Dolby Family Fund (to F.H.G.), AHA-Allen Initiative in Brain Health and Cognitive Impairment award made jointly through the American Heart Association and The Paul G. Allen Frontiers Group 19PABH134610000 (to F.H.G. and C.M.M.); NIH grant R01CA234245 (to C.M.M.), the Lowy Medical Research Institute (to C.M.M.), the Mark Foundation for Cancer Research (to C.M.M.) for metabolic and lipidomic work; R01NS125591 (to J.T.M.) from NINDS for the proteomic work; NIH grants RF1-AG056306 (to J.M. and F.H.G.), P30-AG062429 (to J.M. and F.H.G.), R01-AG085634 (to J.M.); CIRM EDUC4-12804 Interdisciplinary Stem Cell Training Grant (to L.Z.Y.). The NSF-Graduate Research Fellowship Program, the Salk Women and Science Award, the Bert and Ethel Aginsky Research Scholar Award, the Jesse and Caryl Philips Foundation, the Kavli-Helinski Endowed Graduate Fellowship and the Pathways in Biological Science (PiBS) Training Grant (all to J.S.R.).

Thank you to the cores utilized in this study: the Waitt Advanced Biophotonics Core Facility of the Salk Institute (RRID:SCR_014838) with funding from NIH-NCI CCSG P30 CA014195, NIH-NIA San Diego Nathan Shock Center P30 AG068635, The Henry L. Guenther Foundation and the Waitt Foundation, with a special thanks for Sammy Weiser-Novak for the electron microscopy work; the Flow Cytometry Core Facility of the Salk Institute (RRID:SCR_014839) with funding from NIH-NCI CCSG P30 CA014195, and Shared Instrumentation Grants S10-OD023689 (Aria Fusion cell sorter), and S10 OD034268 (Thermo Fisher Bigfoot), with a special thanks for Carolyn O’Connor for senescent cell isolation planning; the Cell Technologies and Engineering Core of the Salk Institute (RRID:SCR_014850) with funding from the NIH-NIA San Diego Nathan Shock Center P30 AG068635, the NIH-NIA Alzheimer’s Disease Research Center P30 AG062429, the AHA Allen Initiative, the California Institute for Regenerative Medicine and the Helmsley Charitable Trust; and the UCI-ADRC with funding from NIH/NIA Grant P30 AG066519. This publication includes data generated at the UC San Diego IGM Genomics Center utilizing an Illumina NovaSeq X Plus that was purchased with funding from a National Institutes of Health SIG grant (#S10 OD026929), with a special thanks to Kristen Jepsen. Thanks go to the staff at Novogene and Plasmidsaurus for RNA sequencing services. We acknowledge the use of BioRender for generation of illustrative cartoons.

## AUTHOR CONTRIBUTIONS

Conceptualization: J.S.R, K.W.R., J.R.J., and F.H.G.; Methodology: J.S.R., K.W.R., T.S.S., M.S.C., S.S., L.Z.Y., J.P.C, and S.C.S.; Formal Analysis: J.S.R., K.W.R., T.S.S., Q.X., M.S.C., S.S., S.S.D, J.R.J.; Investigation: J.S.R, K.W.R., Q.X., T.S.S., M.S.C., S.S., L.Z.Y., C.K.L., V.N.P., R.S.W., S.S.D, A.K., and J.M.P.; Writing – Original Draft: J.S.R.; Writing – Review & Editing: J.S.R., J.R.J., T.S.S., M.S.C., K.W.R., S.S., J.M., S.S.D., J.T.M., C.M.M., and F.H.G.; Supervision: F.H.G., J.R.J., C.M.M., J.T.M., and J.M.

## SUPPLEMENTAL INFORMATION

Tables available upon request.

**Table S1.** Donor line information. DEGs comparing male and female iMG. Donor lines representation per UMAP clusters.

**Table S2.** Top gene markers from microglia clusters in Figure 1. Gene sets utilized from other studies to create module scores in Figure 1.

**Table S3.** DESeq2 results and groups from likelihood ratio test of pseudobulk data of 22 donor iMG split by APOE haplotype. Wald analysis comparing pseudobulk data of APOE44 to APOE33 donor iMG. COMPASS results comparing APOE44 to APOE33 donor iMG. DEGPattern group genes and functional enrichment results per group. Analysis by APOE haplotype separated by sex included.

**Table S4.** DESeq2 results and functional enrichment results from downsampled APOE33 and APOE44 ROSMAP microglia.

**Table S5.** Lipidomics of APOE33 and APOE44 iMG alone and cocultured with iNs. Lipidomics of iNs cocultured with APOE33 and APOE44 iMG.

**Table S6.** Proteomics data of KOLF2.1J and UCI5 APOE33 and APOE44 iMG.

**Table S7.** Bulk RNAseq and functional enrichment results of APOE33+DMSO, APOE44+DMSO and APOE44+LXR iMG.

## Figures

**Figure S1.**
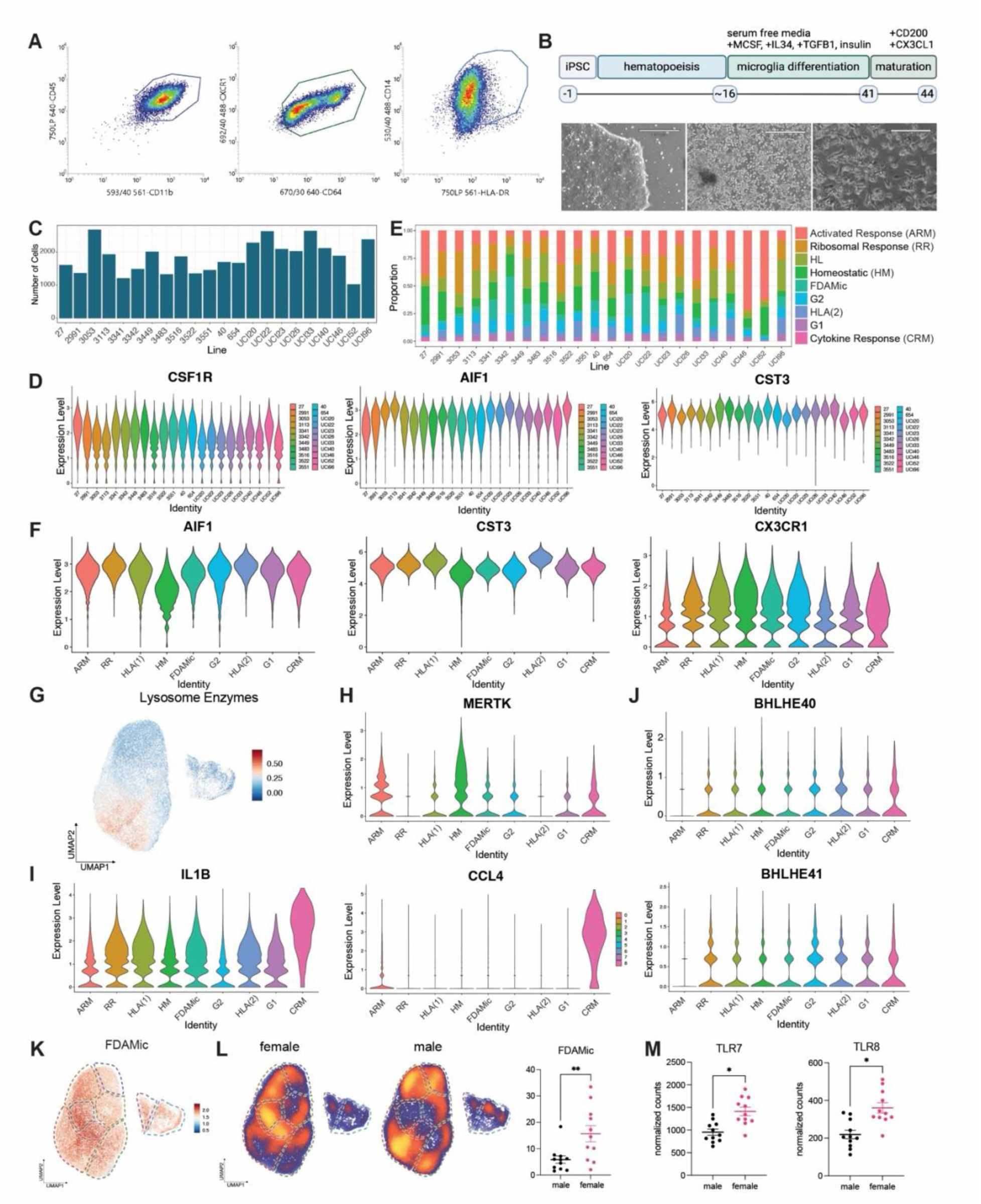
Characterization of iMG and different clusters. (A) FACS gating strategy for collecting iMG positive for CD45, CD11b, CD64, CX3CR1, CD14 and HLA-DRA. (B) Schematic of iPSC conversion to iMG. Left image shows iPSCs, middle image shows iHPCs, and right image shows iMG. Scale bar = 330µm for first two panels, 130µm for last panel. (C) Bar plot represents the number of cells per donor line in the scRNAseq dataset. (D) Violin plots showing expression of microglia-specific genes AIF1, CST3 and CSF1R across the 22 iMG profiled. (E) Bar plot represents the relative contribution of donor lines across clusters. (F) Violin plots showing expression of microglia-specific genes AIF1, CST3 and CSF1R across the 9 identified clusters. (G) Module score of 119 lysosomal enzymes (https://www.proteostasisconsortium.com/pn-annotation/) plotted onto our UMAP. (H-J) Violin plots showing expression of (H) MERTK (I) IL1B, CCL4, and (J) BHLHE40/41 across the different clusters. (K) Module score of FDAMic genes (RPL13, RPLP1, SPP1, RPS19, ACTB, RASGEF1B, PSAP, CSF1R) plotted onto our UMAP. (L) Density plot of UMAP by sex displaying distribution across identified clusters. Proportion of cells in the FDAMic cluster, separated by sex. Each dot represents a donor line. (M) Expression of genes TLR7 and TLR8 across sex. Each dot represents a donor line. (L-M). Unpaired t-test, two-tailed, mean ± SEM *p < 0.05; **p < 0.01

**Figure S2.**
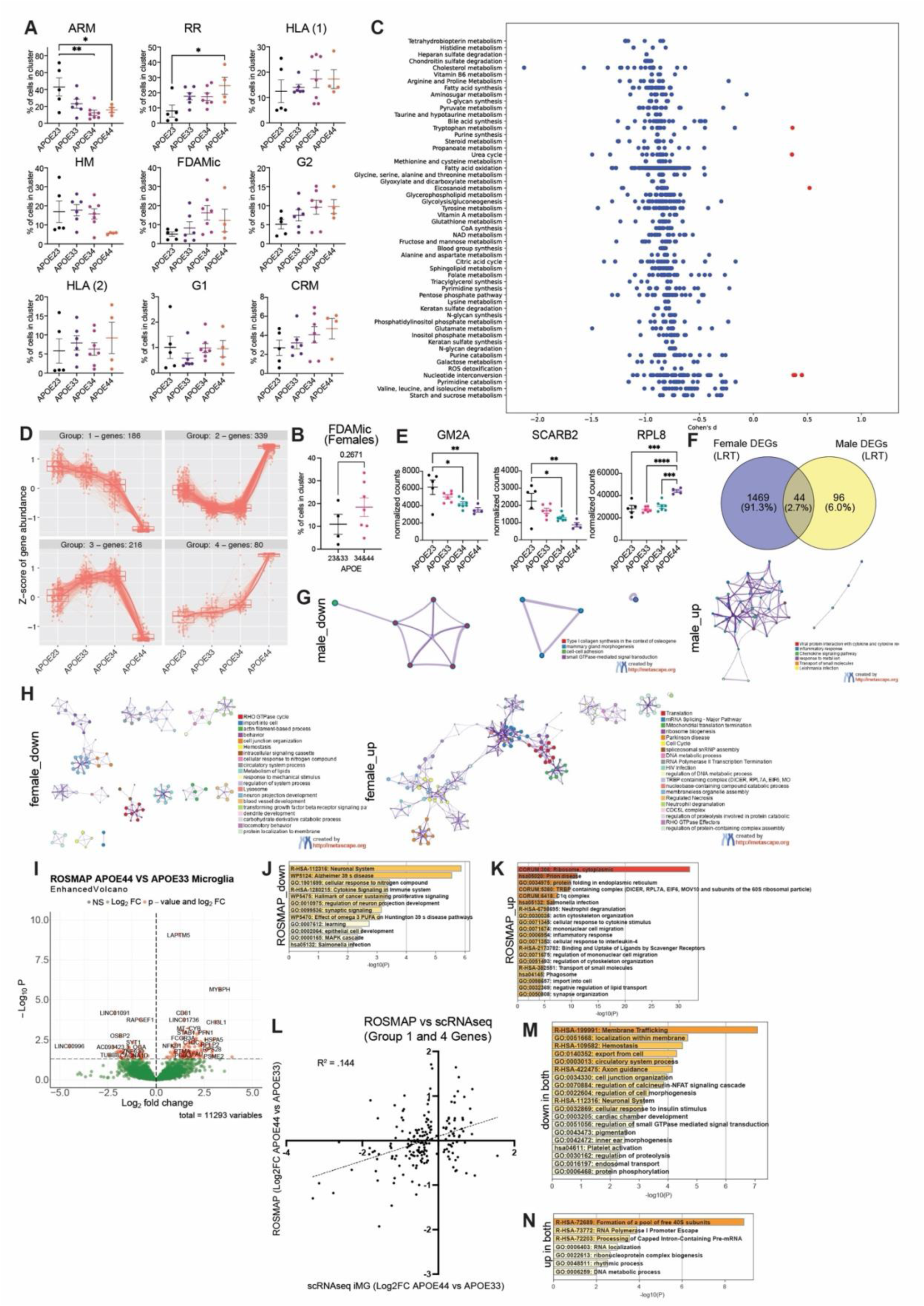
APOE4 iMG have a distinct transcriptomic signature. (A) Distribution and proportion of cells across different cell states, separated by APOE genotype. Each dot represents an individual line. Unpaired t-test, two-tailed, mean ± SEM. (B) Proportion of female cells across FDAMic cluster, separated by APOE4- and APOE4+. Each dot represents an individual line. One way ANOVA, mean ± SEM. (C) Compass analysis of metabolic pathways in APOE44 vs APOE33 lines in G0 phase cells. Wilcoxon signed-rank test, significance set to padj < 0.05 (D) DEGs across APOE haplotypes reveal 4 specific patterns of expression. (E) Expression of genes GM2A, SCARB2 and RPL8 across APOE haplotypes. Each dot represents an individual line. One way ANOVA, mean ± SEM. (F) Venn diagram of number of DEGs with Likelihood Ratio Test (padj < 0.05) separated by sex and analyzed by APOE haplotype. (G,H) Enrichment analyses of downregulated and upregulated genes in (G) male and (H) female lines across APOE haplotype. Significance set to Wald |Log2FC(APOE44 vs APOE33)| > 0, LRT padj < 0.05. Created using Metascape^97^. (I) Volcano plots of ROSMAP APOE44 microglia vs APOE33 microglia. Down sampled to n = 8 for each group. Likelihood Ratio Test, p-values were corrected using the Benjamini-Hochberg correction for multiple testing, significance set to padj < 0.05, |Log2FC cutoff| > 0. (J,K) Enrichment analyses of (J) downregulated and (K) upregulated genes comparing ROSMAP APOE44 microglia vs APOE33 microglia. Significance set to p < 0.05, |Log2FC cutoff| > 0. Created using Metascape^97^. (L) Log2FC changes of genes comparing APOE44 to APOE33 ROSMAP microglia against APOE44 and APOE33 donor iMG. Genes with padj (LRT) < 0.05 for the donor iMG are shown. Log2FCs from Wald analysis are shown for donor iMG. (M-N) Enrichment analyses of (M) downregulated genes in both datasets and (N) upregulated genes in both datasets. Significance set to padj (LRT) < 0.05 for donor iMG. |Log2FC cutoff| > 0 in ROSMAP and donor iMG dataset. Created using Metascape^97^. *p < 0.05; **p < 0.01; ***p < 0.001; ****p < 0.0001

**Figure S3.**
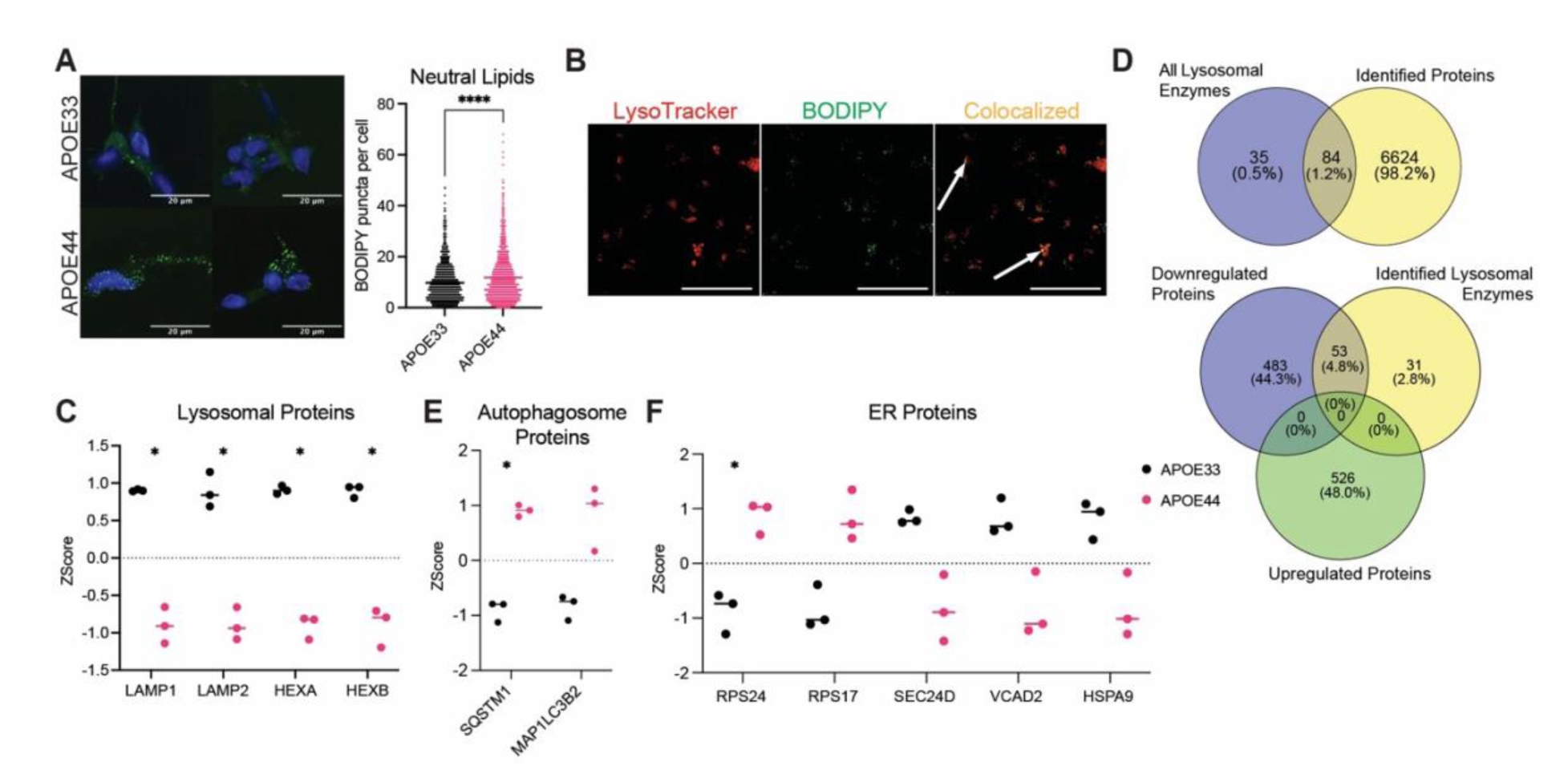
APOE4 iMG have an increase in cholesterol esters in the lysosomes. Proteomics data supports lysosomal and ER stress. (A) Representative images of APOE33 and APOE44 iMG stained for BODIPY-493/503. Scale bar = 20µm. Each dot represents number of puncta per cell (for each group, 3 wells were imaged, 9 frames each). Unpaired t-test, two-tailed, mean ± SEM. (B) BODIPY-493/503, LysoTracker Red DND-99 and colocalization representative image. Scale bar = 50µm. (C) LAMP1, LAMP2, HEXA and HEXB protein levels across APOE33 and APOE44 iMG. (D) Top: Venn diagram shows number of identified lysosomal enzymes in our proteomic analysis (https://www.proteostasisconsortium.com/pn-annotation/). Bottom: Venn diagram shows overlap of significantly downregulated and upregulated proteins with identified lysosomal enzymes. (E) SQSTM1 (p62) and MAP1LC3B2 protein levels across APOE33 and APOE44 iMG. (F) RPS24, RPS17, SEC24D, VDAC2, and HSPA9 protein levels across APOE33 and APOE44 iMG. (C, E, F) Unpaired t-test, two-tailed, mean ± SEM, corrected with Benjamini-Hochberg (n = 3 wells each). *p < 0.05; ****p < 0.0001

**Figure S4.**
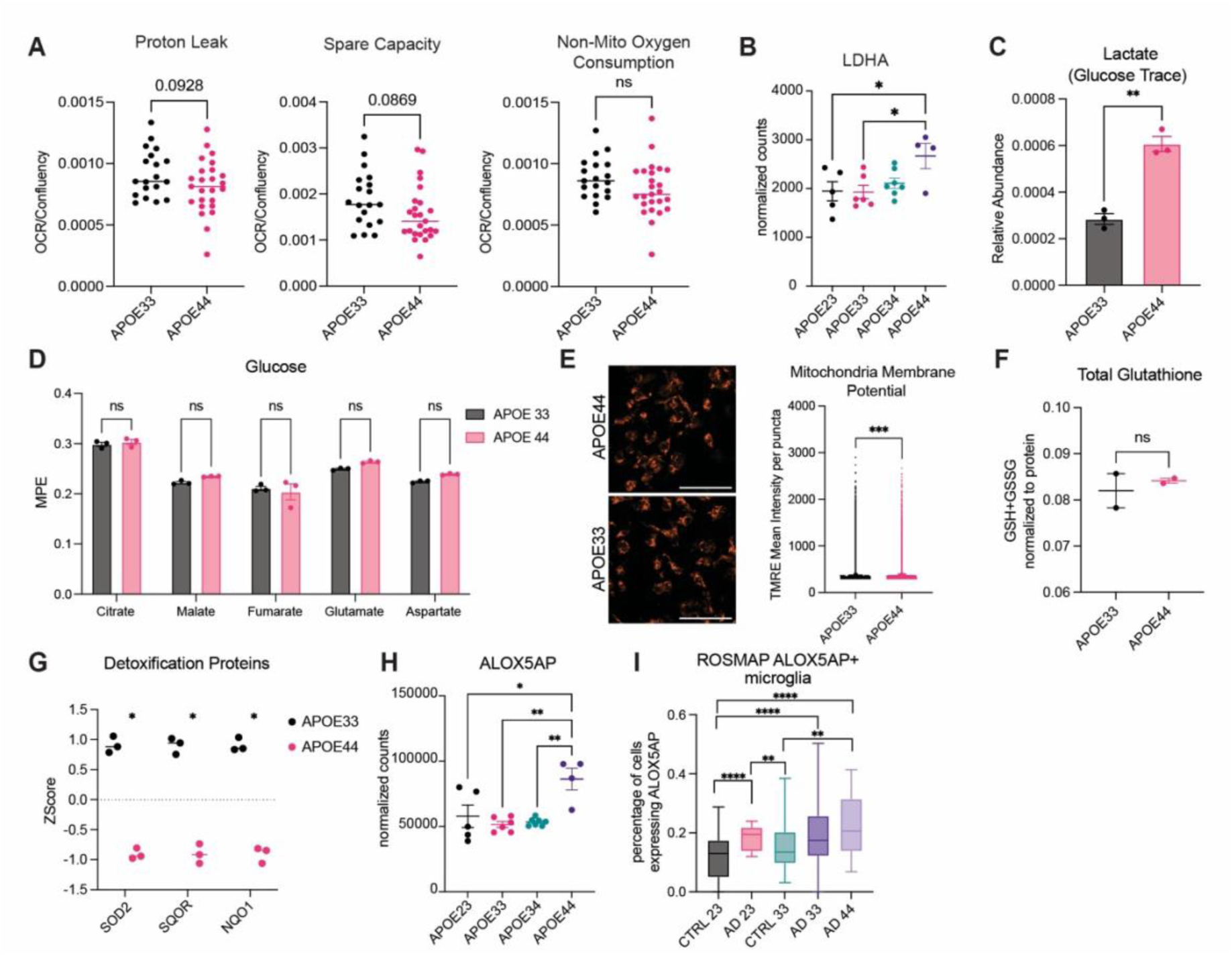
APOE44 iMG exhibit a shift in bioenergetics. (A) Seahorse Mito Stress Test kit. Proton leak, spare capacity and non-mito oxygen consumption normalized to confluency of wells, comparing APOE33 and APOE44 iMG. Each dot represents a replicate well. (B) Expression of LDHA across different APOE haplotypes in the donor iMG dataset. Each dot represents an individual line. One way ANOVA, mean ± SEM. (C) Relative abundance of lactate after tracing with [U-^13^C_6_] glucose. (D) Labeled citrate, malate, glutamate, and aspartate after [U-^13^C_6_] glucose tracing. Two-way ANOVA, Šídák’s multiple comparisons test, mean ± SEM. (E) TMRE staining and quantification of mean intensity per puncta across APOE33 and APOE44 iMG (n = 3 well x 10 frames per line). Scale bar = 50µm. (F) Total glutathione measured from cell lysates using kit (MedChem, HY-K0311). (G) SOD2, SQOR, and NQO1 protein levels across APOE33 and APOE44 iMG. Unpaired t-test, two-tailed, mean ± SEM, corrected with Benjamini-Hochberg (n = 3 wells each). (H) Expression of ALOX5AP across different APOE haplotypes in the donor iMG dataset. Each dot represents an individual line. One way ANOVA, mean ± SEM. (I) Percentage of ALOX5AP+ microglia in the ROSMAP dataset across different APOE haplotypes and diagnoses. Chi-squared test (CTRL APOE23 n = 13, AD APOE23 n = 6, CTRL APOE33 n = 50, AD APOE33 n = 48, AD APOE44 n = 6). (A, C, E-F) Unpaired t-test, two-tailed, mean ± SEM. *p < 0.05; **p < 0.01; ***p < 0.001; ****p < 0.0001. ns, non-significant

**Figure S5.**
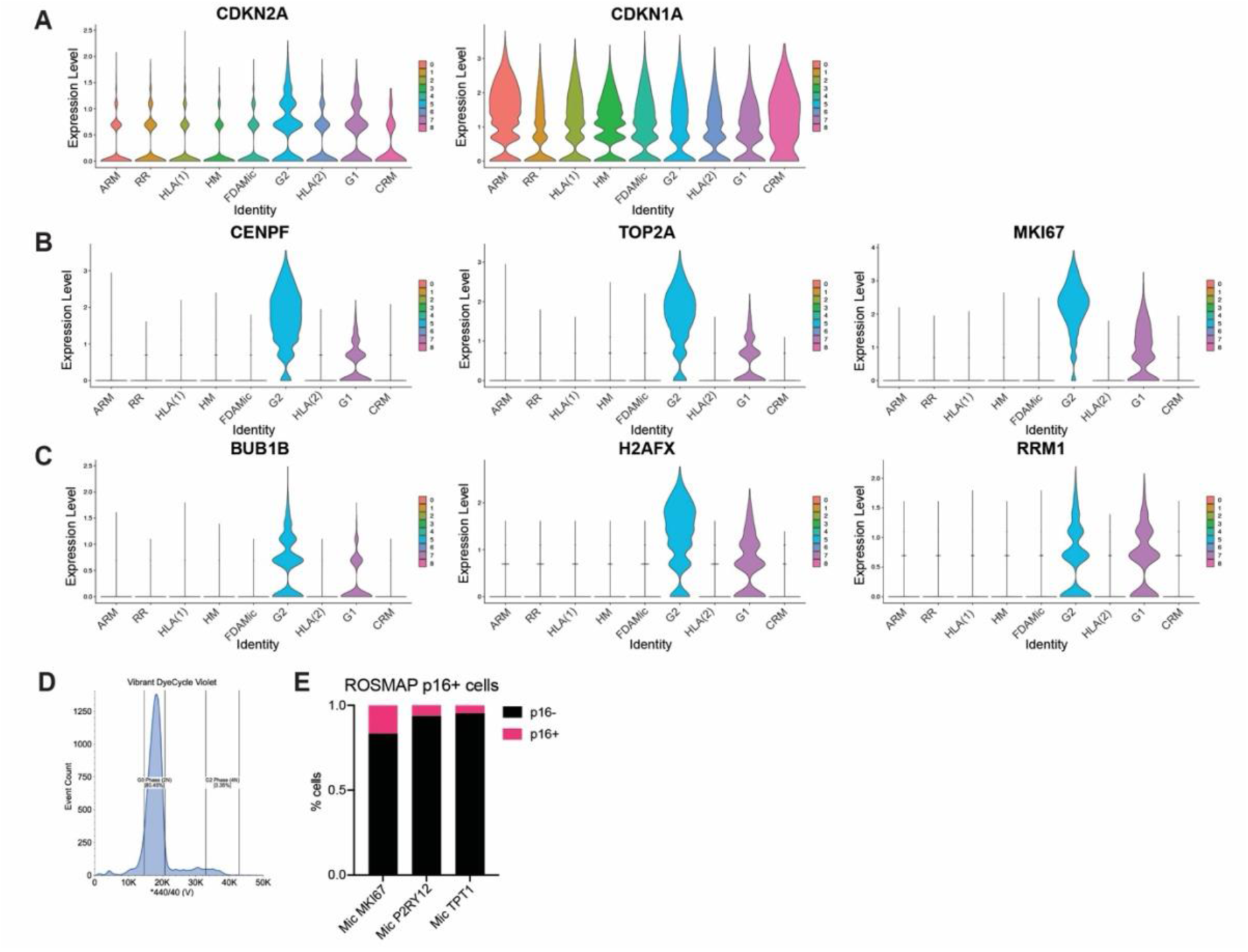
APOE4 promotes a G2 senescent-like state. (A) Violin plots showing expression of senescent markers CDKN2A (p16) and CDKN1A (p21) across the different clusters in the scRNAseq of the 22 donor iMG. (B) Violin plots showing expression of G2 markers and (C) DNA damage markers across the different clusters in the scRNAseq of the 22 donor iMG. (D) FACS gating strategy for sorting out tetraploid (4N) cells using Vibrant DyeCycle Violet and verapamil. (E) Percentage of p16+ microglia in the ROSMAP dataset across the different clusters previously identified.

**Figure S6.**
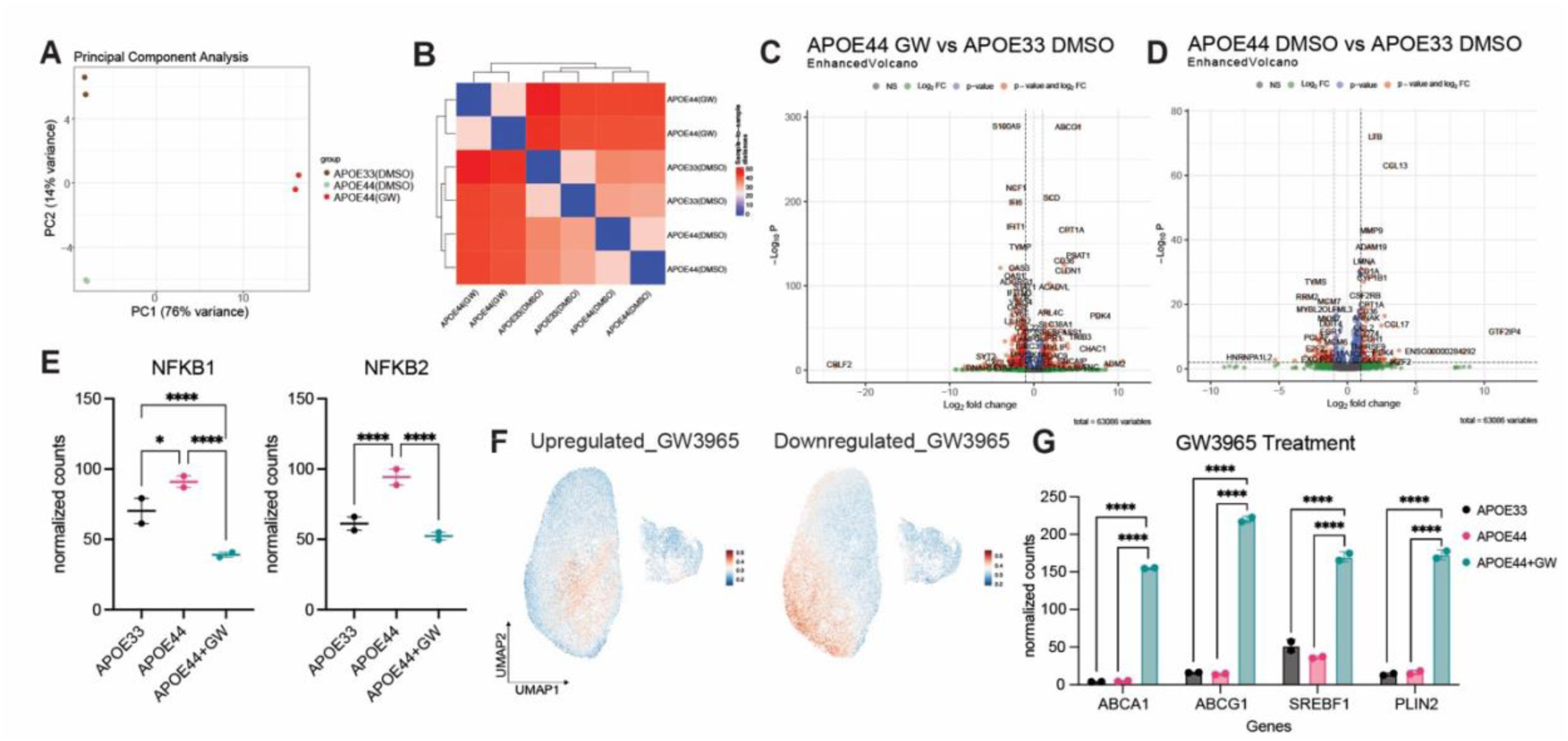
Characterization of LXR agonist GW3965 on APOE44 iMG. (A) Principal component analysis of APOE33(DMSO), APOE44(DMSO) and APOE44(GW) iMG bulk RNAseq (n = 2 wells per group). (B) Euclidean distance analysis of bulk RNAseq comparing APOE33(DMSO), APOE44(DMSO) and APOE44(GW) iMG. (C) Volcano plot of bulk RNAseq comparing APOE44(GW) iMG vs APOE33(DMSO) iMG. Wald test, p-values were corrected using the Benjamini-Hochberg correction for multiple testing, significance set to padj < 0.01, |Log2FC cutoff| > 1. (D) Volcano plot of bulk RNAseq comparing APOE44(DMSO) iMG vs APOE33(DMSO) iMG. Wald test, p-values were corrected using the Benjamini-Hochberg correction for multiple testing, significance set to padj < 0.01, |Log2FC cutoff| > 1. (E) NFKB1 and NFKB2 mRNA expression across groups. Wald test, corrected using the Benjamini-Hochberg correction for multiple testing, mean ± SEM. (F) Module scores created using upregulated genes (left) and downregulated genes comparing APOE44(GW) vs APOE44(DMSO) iMG plotted on our UMAP space. Significance set to padj < 0.01, |Log2FC cutoff| > 1. (G) Expression levels of select genes (ABCA1, ABCG1, SREBF1, PLIN2) across different treatment groups. Each dot represents a duplicate. Wald test, corrected using the Benjamini-Hochberg correction for multiple testing, mean ± SEM.. *p < 0.05; ****p < 0.0001

**Figure S7.**
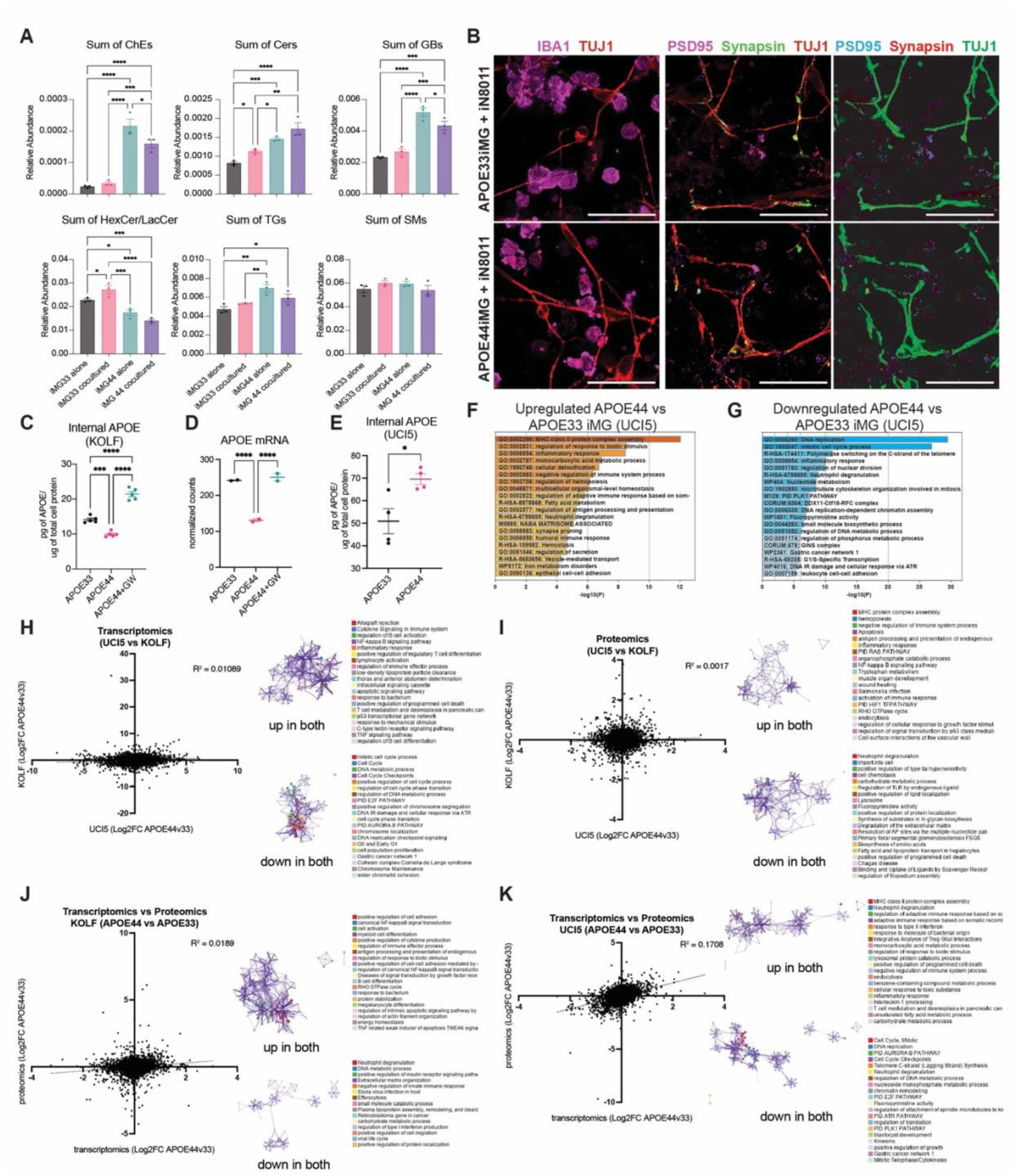
Characterization of iNs and two isogenic APOE33 and APOE44 iMG lines. (A) Sum of lipid species separated by class, plotted across APOE33 and APOE44 iMG cocultured with iNs or alone (n = 3 wells each). (B) Staining of Iba1 and Tuj1 (left); Tuj1, PSD95 and Synapsin (middle) and segmentation (right) in APOE33 iMG + iN8011 (top) and APOE44 iMG + iN8011 (bottom). Scale bar = 50µm. (C) APOE ELISA of cell lysates from KOLF APOE33(DMSO), APOE44(DMSO) and APOE44(GW) normalized to total cell lysate protein (n = 5 wells each). (D) APOE mRNA levels across KOLF APOE33(DMSO), APOE44(DMSO) and APOE44(GW) (n = 2 wells each). Wald test, corrected using the Benjamini-Hochberg correction for multiple testing, mean ± SEM. (E) APOE ELISA of cell lysates from UCI5 APOE33 and APOE44 iMG normalized to total cell lysate protein (n = 4 wells each). Unpaired t-test, two-tailed, mean ± SEM. (F-G) Enrichment analyses of (F) upregulated and (G) downregulated proteins in UCI5 APOE44 vs APOE33 iMG proteomic dataset (n = 3 wells each). Created using Metascape^97^. Unpaired t-test, two-tailed, mean ± SEM, corrected with Benjamini-Hochberg. Significance set to padj < 0.05, |Log2FC cutoff| > 0.5. (H) Gene expression Log2FC for APOE44 vs APOE33 in KOLF iMG compared with corresponding changes in UCI5 iMG. Functional enrichment of overlapping genes. (I) Protein expression Log2FC for APOE44 vs APOE33 in KOLF iMG compared with corresponding changes in UCI5 iMG. Functional enrichment of overlapping genes. (J) Transcriptomic and proteomic Log2FC comparison of APOE44 vs APOE33 in KOLF iMG. Functional enrichment of overlapping genes. (K) Transcriptomic and proteomic Log2FC comparison of APOE44 vs APOE33 in UCI5 iMG. Functional enrichment of overlapping genes. (A, C) One way ANOVA, mean ± SEM. (H-K) Enrichment analyses of overlapping upregulated and downregulated proteins. Significance set to padj < 0.05. Created using Metascape^97^. *p < 0.05; **p < 0.01; ***p < 0.001; ****p < 0.0001

## METHODS

## KEY RESOURCES TABLE

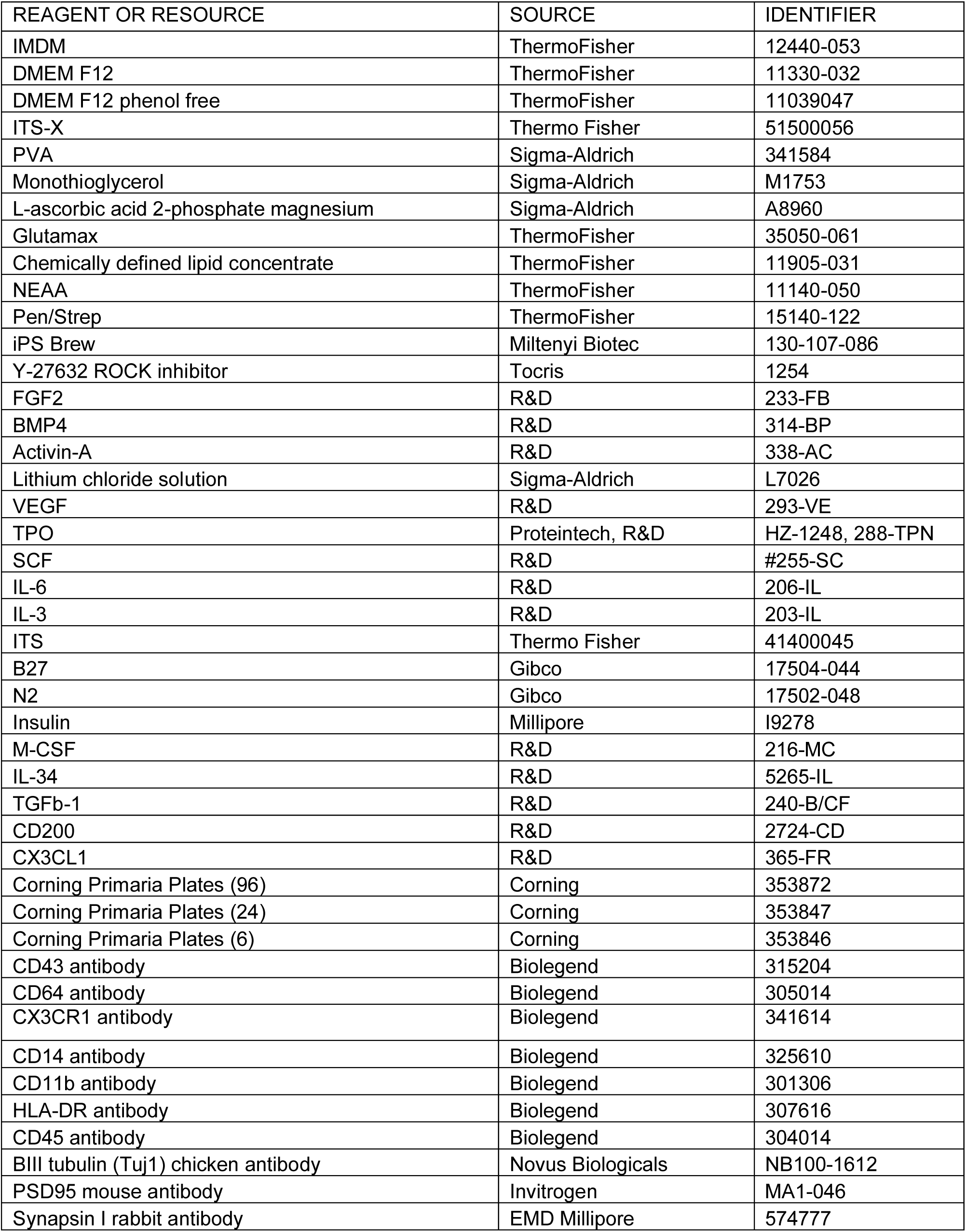
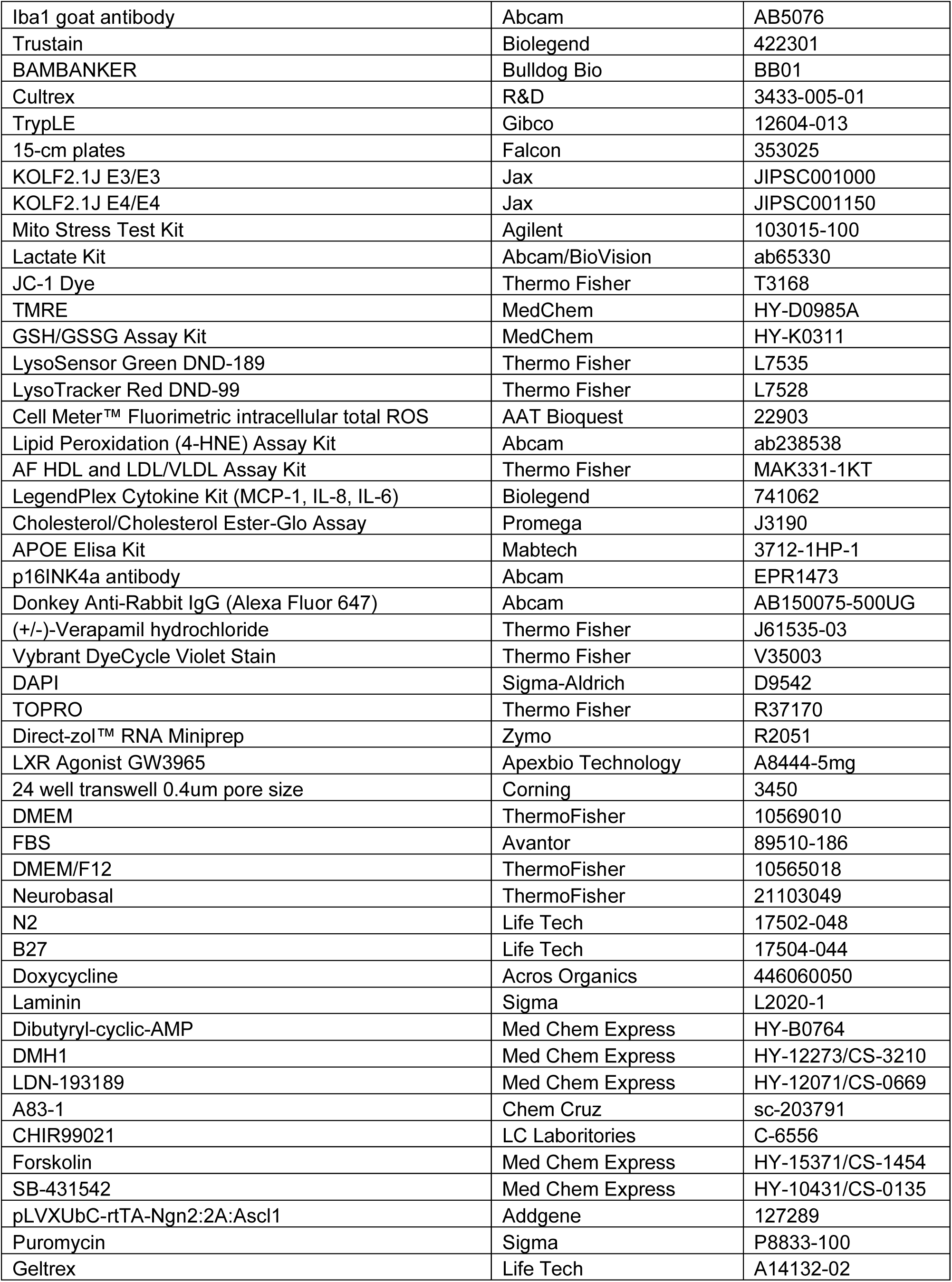
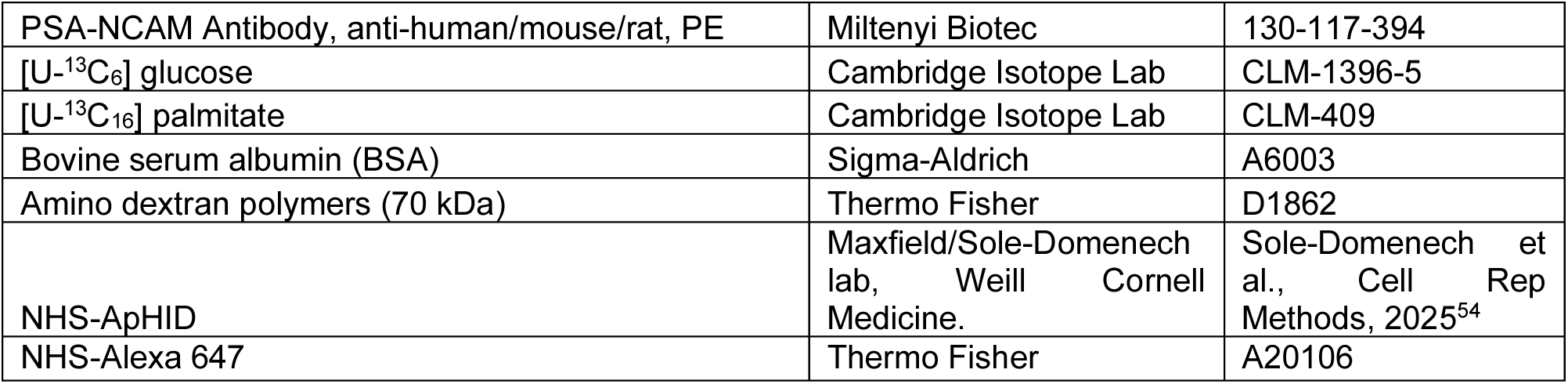

## EXPERIMENTAL MODEL AND STUDY PARTICIPANT DETAILS

The protocol for the use of human induced pluripotent stem cell (iPSC) lines for this study was approved by Salk Institute’s IRB Committee (IRB 09-0003) and the Embryonic Stem Cell Research and Oversight Committee. The Salk Institute is committed to protecting the rights and welfare of human research participants and ensures compliance with all applicable ethical and legal requirements. The iPSC lines used in this study were reprogrammed in two facilities (Salk Institute for Biological Studies, Cell Technologies and Engineering Core; University of California Irvine Alzheimer’s Disease Research Center (ADRC) Induced Pluripotent Stem Cell Core and the Institute for Memory Impairments and Neurological Disorders). iPSCs were maintained in 6-well plates (Corning) in feeder-free conditions using Cultrex (R&D) in iPS Brew (Miltenyi) in a humidified incubator (5% CO2, 37°C). iPSCs were fed fresh media every other day and passaged every 7–8 days.

## METHOD DETAILS

### iMG conversion from iPSCs and maintenance

Using an established protocol with minor modifications, iPSCs were converted into iMG^27^. iPSCs were first induced into hematopoietic progenitor cells (iHPCs) and then into microglia. iPSCs were dissociated using TrypLE (Invitrogen) and plated in a 15-cm tissue culture-treated plate (Falcon) at a density of ∼12million (full 6-well plate 90% confluent) cells per plate with 10 μM ROCK inhibitor (Tocris). The next day (day 0), cells were changed to hematopoietic differentiation media supplemented with FGF2 (50 ng mL−1, R&D), BMP4 (50 ng mL−1, R&D) Activin A (12.5 ng mL−1, R&D) ROCK inhibitor (1 μM, Tocris) and LiCl (2 mM, Sigma) and grown under hypoxic conditions (5% CO2, 5% O2). Hematopoietic differentiation media consisted of IMDM (50%, Thermo Fisher Scientific), DMEM/F12 (50%, Thermo Fisher Scientific), ITSG-X, (2% v/v, Thermo Fisher Scientific), L-ascorbic acid 2-Phosphate magnesium (64 mg mL−1; Sigma), monothioglycerol (400 mM, Sigma), PVA (10 mg mL−1; Sigma), Glutamax (1X, Thermo Fisher Scientific), chemically defined lipid concentrate (1X, Thermo Fisher Scientific), non-essential amino acids (NEAA, 1X, Thermo Fisher Scientific), Penicillin/Streptomycin (P/S Thermo Fisher Scientific, 1% V/V). On day 2, floating cells were collected, spun down, and media was replaced with hematopoietic differentiation media supplemented with FGF2 (50 ng mL−1, R&D) and VEGF (50 ng mL−1, R&D) and returned to hypoxic conditions in the same plate. Day 2 media was equilibrated in hypoxia before use for at least 45 min. On day 4, cells were collected and placed into normoxic conditions (5% CO2, 21% O2) and kept in hematopoietic differentiation media supplemented with FGF2 (50 ng mL−1, R&D), VEGF (50 ng mL−1, R&D) TPO (50 ng mL−1, Proteintech and R&D), SCF (10 ng mL−1, R&D), IL-6 (50 ng mL−1, R&D) and IL-3 (10 ng mL−1, R&D) until colonies released iHPCs (usually between day 14 and 20). iHPCs were then isolated using Fluorescence-activated cell sorting (FACS) by gating on CD43-FITC (Biolegend, 315204)-positive cells. iHPCs were then cultured in microglia differentiation media supplemented with 25 ng ml−1 M-CSF (R&D), 100 ng mL−1 IL34 (R&D) and 50 ng mL−1 TGFβ1 (R&D) for 4 weeks. 100ng mL-1 CD200 (R&D) and 100ng mL-1 (R&D) CX3CL1 were added for 3 days before iMG collection, which occurred after ∼2-4 weeks, and continued during experiments; this media will be referred to as complete microglia media. Microglia differentiation media consisted of DMEM/F12, ITS-G (2%v/v,Thermo Fisher Scientific), B27 (2% v/v), N2 (0.5%, v/v), monothioglycerol (200 μM), Glutamax (1X), NEAA (1X), and insulin (5 μg mL-1; Sigma). iMG were FAC sorted for microglia markers CD64, CX3CR1, CD14, CD11B, HLADR, and CD45 and plated for subsequent experiments or collected immediately for scRNAseq.

### Sequencing collection

To isolate for scRNAseq, iMG were sorted for microglia markers CD64, CX3CR1, CD14, CD11B, HLADR, and CD45 and subsequently resuspended in BSA 0.04%. scRNAseq was conducted using 10X Chromium Next GEM Single Cell 5’ Kit v2 (10x, PN-1000263). Cells from multiple lines were pooled in equal proportions, from which 40,000 cells were loaded per 10x reaction and computationally demultiplexed using *demuxlet*^98^. Droplet emulsions for the reverse transcription reaction were created using the 10X Genomics Chromium controller. Each reaction’s library was sequenced with ∼400 million, 101bp paired-end reads on NovaSeq S4 flowcells.

### Sample genotyping

Genomic DNA from each iPSC line was submitted for genotyping on an Illumina Infinium CoreExome-24 (v1.4) chip. Genotyped variants were called against hg19 using the IlluminaGenotypingArray_v1.12.10 WDL analysis pipeline (github.com/broadinstitute/warp) and lifted over to hg38 using GATK LiftOverVcf. Variant imputation was conducting on the Michigan imputation server using all populations from the 1000g-phase-3-30x reference panel (https://imputationserver.sph.umich.edu).

### scRNA-seq Quality control

Sequencing reads were mapped to hg38 and gene expression counts were quantified using the STARsolo command from STAR v2.7.11b with flags to handle 5’ v2 chemistry (--clip5pNbases 39 0 --soloType CB_UMI_Simple --soloCBstart 1 --soloUMIstart 17 --soloCBlen 16 --soloUMIlen 10), rescue multimapping reads (--outFilterMultimapNmax 100 --winAnchorMultimapNmax 100 --soloMultiMappers EM), to map the additional bases of the barcode read (--soloBarcodeReadLength 101), to use intronic reads in quantification (--soloFeatures GeneFull), and to mimic CellRanger in empty droplet filtering (--soloCellFilter EmptyDrops_CR). Cells with nCount_RNA < 2000 or more than 15% mitochondrial reads were discarded. *Demuxlet* was used with imputed variant calls from genotyping to match scRNA profiles to iMG lines. Doublets were removed using *scrublet* and *demuxlet*. SCTransform was used to normalize counts for PCA and batch correction. For all other downstream analysis, counts were CPM scaled and log-normalized.

### Microglia clustering

We performed integration across sequencing batches using SCTransform. Using the Seurat package, we used the first 15 principal components based on the top 2,000 highly variable genes to conduct Louvain clustering and UMAP visualization. We created module scores with the AddModuleScore function using existing gene signature sets from other microglia studies (Table 2) to assist in characterizing each of these clusters. Lysosomal enzyme genes were found on the Proteostasis Consortium database under “Lysosome Catabolism” (https://www.proteostasisconsortium.com). The R package LSD was used to plot and compare sample distribution in UMAP space. FindAllMarkers R function was use to find top markers from each cluster (min.pct = 0.6, logfc.threshold = 0.5). Top 5 markers were plotted against each cluster using DoHeatmap R function, with normalized expression scaled by gene.

### Microglia pseudobulk differential gene expression

The counts per individual from cells considered in the G0 phase were summed up to create the pseudobulk counts. We used DESeq2 to normalize and perform the differential analysis. We followed standard analysis steps with standard parameters. We used the Likelihood Ratio Test, which utilizes Analysis of Deviance (ANODEV), to compare different APOE haplotypes (APOE23, APOE33, APOE34, APOE44) reduced by sex. Four groups were created that revealed groups of cells with similar expression patterns. DEGs were defined as those with an adjusted p-value < 0.05. All enrichment analyses were done using Metascape^97^.

### iMG G2 (4N DNA) isolation

To collect cells with 4N DNA content (G2) and 2N DNA content (G0), iMG were treated with verapamil (50µM) to block efflux of Vybrant DyeCycle Violet (DCV; Thermo V35003) for 30 min at 37 C at a concentration of 1×10^6^ cells mL-1. Subsequently, without washing the verapamil out, DCV was added at a concentration of 5µM for another 30 min at 37 C. Cells were sorted at room temperature immediately after treatment. Gates were created to collect 2N and 4N DNA content and plated at a density of 25K cells per well in a 96-well plate for subsequent experiments, or 100K per well in a 24-well plate for lipid extraction.

### Cell arrest assay

To observe if 4N cells were arrested, after 72 hours of plating, verapamil and DCV were used again, and fluorescence of DCV was measured using a plate reader with an excitation of 405 nm and emission of 440 nm. Fluorescence was normalized to cell count, as measured by Incucyte.

### JC-1

Cells were plated at a density of 50K cells/well in a black-walled 96-well plate. To measure MMP, JC-1 dye was used at a concentration of 2µM in phenol free DMEM/F12 for 30 min and washed off (5x half media washes) before measuring on a plate reader (Glomax). For the green fluorescence, the plate was read with an excitation of 480 nm and emission of 535 nm. For the red fluorescence, the plate was read with an excitation of 550 nm and emission of 600 nm. The dye exists as a monomer at low concentrations and exhibits an emission at ∼535 nm, and at higher concentrations, the dye forms aggregates, which are found in healthy mitochondria, that exhibit an emission maximum at ∼590 nm. To calculate MMP, we took the ratio of red fluorescence (aggregates) to the green fluorescence (monomers).

### Seahorse Mito Stress Test Kit

Seahorse XFe96 Mito Stress Test Kit was utilized using manufacturer’s instructions. A total of 50K cells from APOE33 and APOE44 isogenic iMG were plated into the provided 96-well plate, with 32 wells each. Pyruvate (1 mM), glutamine (2 mM) and glucose (10 mM) were added to DMEM/F12 no glucose, no glutamine (Biowest, L0091) to serve as base media. Oligomycin (1.5 µM) was delivered in Port A, Carbonyl cyanide-4 (trifluoromethoxy) phenylhydrazone (FCCP) (1 µM) in Port B, and Rotenone/Antimycin (0.5 µM) in Port C. Baseline readings were measured for 8 cycles, with 3-min mixes and 3-min measurements. Readings after drug delivery were measured for 3 cycles, with 3-min mixes and 3-min measurements. Images of each well were taken on Incucyte and cells were counted in ImageJ for normalization of data. During analysis, wells that exhibited failed drug delivery were excluded from analysis.

### ROS

Cells were plated at a density of 25K cells/well in a black walled 96-well plate. Media was replaced with 100ul DMEM/F12 phenol free (Thermo) and ROS Brite 670 following manufacturer’s instructions. The plate was imaged on Glomax reader at an excitation of 627nm and emission of 660-720nm after 30 min and normalized to cell count, which was quantified on ImageJ after taking images of the wells on Incucyte.

### HNE

Cells were plated at a density of 50K cells/well in a black-walled 96-well plate. The Lipid Peroxidation (4-HNE) Assay Kit (Abcam) was used following manufacturer’s instructions and normalized to protein amount, as quantified by Pierce BCA Protein Assay Kit (Thermo, 23225/23227/A65453).

### Cytokine assay

Cells were plated at a density of 250K cells/well in a 12-well-plate. Media was replaced with 500ul complete microglia media and collected after 24 hours. Supernatant was centrifuged to get rid of debris and immediately assayed with LegendPlex (IL6, CCL2, IL8 – Biolegend #741062 mix and match kit) using manufacturer instructions and normalized to cell count which was quantified on ImageJ after taking images of the wells on Incucyte immediately before collecting media.

### Ratiometric lysosomal pH measurements

pH imaging using ApHID. iMG were plated at a density of 70K cells/well in a glass bottom black-walled 96-well plate (Cellvis, P96-1-N) coated with Cultrex in 100µl phenol free complete microglia media with or without organoid debris. After 48 hours, 70 kDa amino dextrans labeled with the pH sensor NHS-ApHID^54^ and NHS-Alexa 647 (pH-independent, Thermo, A20106) were added to the cell media at a concentration of 0.5 mg/mL and incubated overnight. Next morning, cells were carefully washed in fresh medium (5 half media washes to prevent cell detachment) and chased for 4h to ensure late endosomal and lysosomal dextran localization. Organoid debris was maintained in the treated groups. 3 wells were fixed in 0.5% paraformaldehyde (prepared in 1X PBS) for 5 min at 37 °C followed by 5 washes in 1X PBS and reserved for use as calibrators. The plate was transferred to a fluorescence imaging enclosure at 37 °C supplied with 5% CO2 and allowed to equilibrate for 20 min followed by imaging. Fixed cells were incubated in 50 mM TRIS maleate buffer containing 40 mM methylamine hydrochloride, 40 mM sodium acetate and 40 µM monensin as membrane-permeant equilibrators, adjusted to pH 5.0, for 30 at 37 °C and imaged. ApHID and Alexa 647 were excited with 495 nm and 645 nm laser lines, and emitted light was collected in the spectral window of 500-550 nm and 660-720 nm, respectively. Stacks of images containing both green and far-red channels were acquired using a 20X air objective, and background signal was eliminated by removing the 10th signal percentile for each frame. Alexa 647 signal was selected by applying a signal threshold. A binary mask was generated and applied to the ApHID and Alexa 647 channels, corresponding to late endosomal and lysosomal compartments. Filtered ApHID and Alexa 647 signal was measured for each frame and ApHID/Alexa 647 fluorescence ratios calculated using FIJI and Metamorph digital image analysis software. Fluorescence ratios measured in fixed cells at pH 5.0 were calculated and used to construct a ratio-to-pH calibration using titration data for the same dextrans acquired in solution (see Sole-Domenech et al., Cell Rep Methods, 2025). The resulting ratio-to-pH calibrations were fit to 4-component sigmoidal curves and used to interpolate fluorescence ratios, measured in live cells, to their corresponding pH values.

### Digital Image Analysis

Digital image analysis and quantification was done using FIJI (ImageJ) 4 version 1.54f for Windows (https://fiji.sc/) and MetaMorph version 6.7.1 for Windows (Molecular Devices, San Jose, California USA, www.moleculardevices.com).

### Organoid debris extraction

Human iPSC-derived glial-enriched organoids were generated using an established protocol and maintained in culture for 2 years^99^. To prepare organoid debris, organoids were lysed in 500 µL of M-PER Mammalian Protein Extraction Reagent (Thermo, 78501) according to the manufacturer’s instructions. Lysates were centrifuged to separate soluble proteins from cellular debris, and the resulting debris pellet was resuspended in complete microglia media for treatment of iMG in lysosomal acidity assays.

### DQ-BSA

To assess lysosomal degradation capacity, a DQ-BSA assay was performed. iMG were seeded at 50K cells per well in a glass-bottom 96-well plate (Ibidi, 89627) coated with Cultrex. After 48 hours, cells were incubated with DQ-BSA (MedChem, HY-D2449) at 10 µg/mL in complete microglia medium for 4 hours. Following incubation, DQ-BSA was thoroughly removed with 5 half-volume media washes. For inhibition controls, cells were pretreated with 10 nM bafilomycin prior to DQ-BSA addition. Imaging was performed 16 hours post-treatment, and quantification methods are described in the Confocal Microscopy and Image Analysis section.

### BODIPY and Lysotracker

iMG were seeded at 50K cells per well in a glass-bottom 96-well plate (Ibidi, 89627) coated with Cultrex. To assess neutral lipid accumulation, BODIPY (2 µM) was added to cells and imaged after 30 min. To assess colocalization of neutral lipids in lysosomes, BODIPY (2µM) and Lysotracker DND-99 (100 nM) were added and imaged after 30 min. Imaging and quantification methods are described in the Confocal Microscopy and Image Analysis section.

### TMRE

iMG were seeded at 50K cells per well in a glass-bottom 96-well plate (Ibidi, 89627) coated with Cultrex. To assess MMP, as an orthogonal approach to JC-1, TMRE (50 nM) was added to cells and imaged after 30 min. Imaging and quantification methods are described in the Confocal Microscopy and Image Analysis section.

### Glutathione

iMG were seeded at 100K cells per well in a 96-well plate (Corning Primaria, 353872). GSH/GSSG assay kit was performed using manufacturer instructions. Total levels were normalized to total protein levels of cell lysates.

### Confocal microscopy and imaging analysis

Cells were imaged with Celldiscoverer 7 (Zeiss), with 40x water immersion objective in Airyscan mode. Acquired images were analyzed by Arivis (Zeiss). Cellpose-Based Segmenter (CP model) was used for cell segmentation, which was then subjected to a volume filter (>100 µm³) to remove unhealthy cells and cell debris. Machine Learning Segmenter was used for puncta segmentation, which was then subjected to volume filter (>0.2 µm³) to remove artifacts introduced by the brightness heterogeneity of cell bodies. The numbers of cell count, and punctum count were quantified, as described previously^56^.

### APOE ELISA

Cells were plated at a density of 50K cells/well in a 96-well plate in 100 µL complete microglia media. After 24 hours, 50 µL of complete microglia media was added containing GW3965 (Apexbio Tech, A8444-5mg) or DMSO to bring the final concentration of GW3965 to 10 µM. After an additional 24 hours, supernatant and cells were collected for APOE ELISA analysis (Mabtech, 3712-1HP-1) using manufacturer’s instructions. 75 µL of the supernatant was diluted 1:1 with the APOE ELISA Buffer and 100 µL was used for downstream analysis. Cells were lysed with 100µL M-PER Mammalian Protein Extraction Reagent (Thermo, 78501) and Halt Protease Inhibitor (Thermo, 78440) following manufacturer’s instructions. 75 µL of the cell lysate was diluted 1:1 with the APOE ELISA Buffer and 100 µL was used for downstream analysis. Remaining cell lysate was quantified with Pierce BCA Protein Assay Kit (Thermo, 23225/23227/A65453) for normalization.

### HDL Measurement

Cells were plated at a density of 50K cells/well in a 96-well plate in 100 µL phenol-free complete microglia media. Media was collected for HDL quantification using the Sigma AF HDL and LDL/VLDL Assay Kit (MAK331-1KT), following manufacturer’s instructions and measured on plate reader Glomax. Protein was extracted from cells and the lysate was quantified with Pierce BCA Protein Assay Kit (Thermo, 23225/23227/A65453) for normalization.

### Glucose and Palmitate Tracing

Cells were plated in 24-well plates (Corning, 353847) at a density of 250K cells/well. For glucose tracing, media was replaced with DMEM/F12 (Biowest, L0091), ITS-G (2%v/v,Thermo Fisher Scientific), B27 (2% v/v), N2 (0.5%, v/v), monothioglycerol (200 μM), Glutamax (1X), NEAA (1X), and insulin (5 μg mL-1; Sigma), 25 ng mL-1 M-CSF (R&D), 100 ng mL-1 IL34 (R&D), 50 ng mL-1 TGFβ1 (R&D), 100 ng mL-1 CX3CL1 (R&D), 100 ng mL-1 CD200 (R&D), 15 mM HEPES, 4.5 mM glutamine and 17.5 mM [U-^13^C_6_] glucose (Cambridge Isotope Laboratories, Inc., CLM-1396-5) for 24 hours before extraction. For palmitate tracing, complete microglia media was supplemented with 100 µM [U-^13^C_16_] palmitate (Cambridge Isotope Laboratories, Inc., CLM-409) conjugated to BSA (Sigma-Aldrich, Cat#A6003) for 24 hours.

### Metabolite extraction and GC-MS analysis

At the conclusion of the tracer experiment, media was aspirated. Then cells were rinsed twice with 0.9% saline solution. Five nmol of norvaline were added to each well along with deuterated lipid standards: 0.5 µL of ultimate splash (Avanti Polar Lipids, Cat#330820), 5 nanomoles of cholesterol-d7 (Cambridge Isotope Laboratories, Inc., DLM-3057) 25 pmol d18:1-d7/15:0 glucosylceramide (Avanti Polar Lipids, Cat# 330729), 25 pmol d18:1-d7/15:0 lactosylceramide (Avanti Polar Lipids, Cat# 330727). 500 µL of ice-cold methanol was added to each well, then cells were scraped from the plate and lysates transferred to Eppendorf tubes and sonicated for 5 min. Separation of polar and organic phases was performed by addition of 200 µL of water and 500 μL of chloroform. Samples were then vortex and centrifuged at 21,000 gs for 5 min at 4°C. The upper phase was transferred to a GC vial and dried overnight under vacuum at 4°C. The organic phase was collected and 2 μL of formic acid was added to the remaining polar phase, which was re-extracted with 500 μL of chloroform. Combined organic phases were dried under nitrogen and used for neutral lipid analysis. The dried samples were resuspended in 60 μL of the initial running buffer and then analyzed with LC-MS/MS. 5 μL of sample were injected.

Dried polar metabolites were processed for gas chromatography-mass spectrometry (GC-MS) as described previously by Cordes and Metallo^100^. Briefly, polar metabolites were derivatizedusing a Gerstel MultiPurpose Sampler (MPS 2XL). Methoxime–*tert*-butyldimethylchlorosilane (tBDMS) derivatives were formed by the addition of 15 μL of 2% (w/v) methoxylamine hydrochloride (MP Biomedicals, Solon, OH) in pyridine and incubated at 45°C for 60 min. Samples were then silylated by the addition of 15 μL of *N*-*tert*-butyldimethylsily-*N*-methyltrifluoroacetamide (MTBSTFA) with 1% tBDMS (Regis Technologies, Morton Grove, IL) and incubated at 45°C for 30 min.

Derivatized polar samples were injected into a GC-MS using a DB-35MS column (30 m by 0.25 mm i.d. by 0.25 μm; Agilent J&W Scientific, Santa Clara, CA) installed in an Agilent 7890B GC system integrated with an Agilent 5977a MS. Samples were injected at a GC oven temperature of 100°C that was held for 1 min before ramping to 255°C at 3.5°C/min and then to 320°C at 15°C/min and held for 3 min. Electron impact ionization was performed with the MS scanning over the range of 100 to 650 mass/charge ratio (*m/z*) for polar metabolites. Metabolite levels and mass isotopomer distributions were analyzed with an in-house MATLAB script that integrated the metabolite fragment ions and corrected for natural isotope abundances. Mole percent enrichment (MPE) was calculated *via* the following equation:

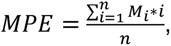, where n is the number of carbons in the molecule and M_i_ is the corresponding isotopologue.

### LC-MS Lipidomic Analysis

Chromatographic separation and lipid species identification for neutral lipids was performed using Q Exactive orbitrap mass spectrometer with a Vanquish Flex Binary UHPLC system (Thermo Scientific) equipped with an Accucore C30, 150 × 2.1 mm, 2.6 µm particle (Thermo) column at 40°C. Chromatography was performed using a gradient of 40:60 v/v water:acetonitrile with 10 mM ammonium formate and 0.1% formic acid (mobile phase A) and 10:90 v/v acetonitrile: propan-2-ol with 10 mM ammonium formate and 0.1% formic acid (mobile phase B), both at a flow rate of 0.2 ml min−1. The liquid chromatography gradient ran from 30% to 43% B from 3-8 min, then from 43% to 50% B from 8-9 min, then 50–90% B from 9–18 min, then 90–99% B from 18–26 min, then held at 99% B from 26–30 min, before returning to 30% B in 6 min and held for a further 4 min. Neutral lipids were analyzed in positive mode using spray voltage 3.2 kV. Sweep gas flow was 1 arbitrary unit, auxiliary gas flow was 2 arbitrary units, and sheath gas flow was 40 arbitrary units, with a capillary temperature of 325°C. Full MS (scan range, 200 to 2000 *m/z*) was used at 70,000 resolution with 10E6 automatic gain control and a maximum injection time of 100 ms. Data-dependent MS2 (Top 6) mode at 17,500 resolution with automatic gain control set at 10E5 with a maximum injection time of 50 ms was used.

Data were analyzed using EI-Maven. Mass error was set to 5 ppm for metabolite identification. Mass isotopomers were filtered to elute within 5 seconds of the unlabeled parent ion. Lipid species-specific fragments used for identification and quantitation are presented in Table S5. Relative abundance was calculated by normalizing to internal standards specific to the lipid class and protein per sample. Absolute abundances were calculated by normalizing to specific lipid species in Ultimate splash or specific deuterated standards added during extraction. Mass isotopologue distributions were analyzed with an in-house MATLAB script that integrated the metabolite fragment ions and corrected for natural isotope abundances.

### Fibroblast to neuron (iN) differentiation

Primary human fibroblasts were originally derived from skin biopsies and maintained in DMEM supplemented with 15% fetal bovine serum and 1% non-essential amino acids. These cells were then transduced using lentivirus carrying the pLVXUbC-rtTA-Ngn2:2A:Ascl1 construct (Addgene: #127289) and subsequently purified with puromycin (Sigma, 1 µg mL-1). Following selection, the cells were pooled at high densities (approximately 3×10⁴ to 5×10⁴ cells/cm²) and transitioned into a neural conversion medium. This medium consisted of a 1:1 mixture of DMEM:F12 and Neurobasal (ThermoFisher), supplemented with N2 and B27 (ThermoFisher), doxycycline (Sigma-Aldrich, 2 µg mL-1), Laminin (Sigma-Aldrich, 1 µg mL-1), dibutyryl-cyclic-AMP (Santa Cruz, 100 µg mL-1), DMH1 (Sanova Pharma, 5 µM), LDN-193189 (Sanova Pharma, 0.5 µM), A83-1 (Santa Cruz, 0.5 µM), CHIR99021 (LC Laboratories, 3 µM), forskolin (LC Laboratories, 5 µM), and SB-431542 (MedChem, 10 µM). The media was changed every other day for a total of 21 days before the cells were sorted using MACS Cell Separation (Miltenyi Biotec) with PSA-NCAM Antibody (Miltenyi Biotec, 130-117-394) and plated for subsequent experiments.

### iN and iMG coculture experiments

For lipid transfer experiments, APOE33 and APOE44 (KOLF) isogenic iMG were plated in 24-well plates (Corning, 353847) at a density of 250K cells/well. iNs were plated in 24 mm, 0.4µm pore Transwells (Corning, 3450) coated with Geltrex (Life Tech) at a density of 90K cells/transwell. In the first experiment, iNs at day 35 were cocultured with iMG for 7 days before extraction in DMEM/F12 (Thermo), 2% B27-vitA, 0.5% N2, 1x NEAA, 1x Glutamax, insulin (5 μg mL-1; Sigma), 2% ITS, 1x BME, 25 ng mL-1 M-CSF (R&D), 100 ng mL-1 IL34 (R&D), 50 ng mL-1 TGFβ1 (R&D), 100ng mL-1 CX3CL1 (R&D), and 100ng mL-1 CD200 (R&D); this media will be referred to as GOSF. Half media changes were done every other day for 7 days. Lipidomic analysis was performed at the end of the experiment.

For microglia-neuron co-culture experiments assessing synaptic effects, neurons were seeded at a density of 133,000 cells per well in 8-well Ibidi chambers coated with Geltrex. After 2 days, microglia were added at a density of 150,000 cells per well. Cultures were maintained in GOSF medium, with half-media changes performed every other day for 7 days prior to fixation. Cells were then fixed and immunostained for Synapsin, PSD95, and Tuj1 to assess synaptic structure, or for Iba1 and Tuj1 to evaluate microglia-neuron interactions. Imaging and quantification methods are described in the Confocal Microscopy and Image Analysis section.

### Proteomics sample preparation

Microglia cells were lysed by resuspending the pellets in 8 M urea solution. The lysates were treated with dithiothreitol (5mM) and incubated at 60°C for 30 min with constant shaking at 500 rpm. Post-reduction, the cysteine residues were alkylated with iodoacetamide (40mM) at 60°C for 30 min. The protein extracts were digested, first with LysC (enzyme: substrate= 1:100) for 2 h at 25°C, and then with trypsin (enzyme: substrate= 1:50) for 3 h at 37°C with shaking at 500 rpm. The digestion reaction was quenched by acidifying the solution to 0.1% formic acid. The digested peptides were desalted with C18 solid-phase extraction, concentrations were determined by BCA, and samples were stored at -20°C until being subjected to liquid chromatography-tandem mass spectrometry (LC-MS/MS).

### LC-MS/MS proteomics data acquisition

Mass spectrometric analysis of peptides derived from the KOLF male line was performed using a QExactive HF-X mass spectrometer (Thermo Scientific, San Jose, CA) outfitted with a Nanospray Flex™ Ion Source (Thermo Scientific, San Jose, CA) ionization interface, and attached to a Thermo Dionex Ultimate 3000 liquid chromatography (LC) platform. Data were acquired for 120 min following a 20.6-min delay from sample injection. The LC platform was configured to load the sample on an on-line trapping column packed in-house (4 cm x 100µm i.d. x 360 µm o.d. fused silica; Polymicro Technologies Inc., Phoenix, AZ), followed by separation on an analytical column (30 cm × 75 μm i.d. x 360 µm o.d., packed with 1.7 μm particle size C18 media; Waters Acquity BEH). Sample injection was performed by loading a 5 µL volume of sample at 7 µL/min for 4 mins unto a trap column. Following the cleanup step, the sample was transferred to the analytical column using reverse-flow elution at 200 nL/min. Mobile phases consisted of buffer A (0.1% formic acid in Water) and buffer B (0.1% formic acid in Acetonitrile) with peptide elution performed using the following gradient profile (min, %B): 0.0, 1.0; 12.6, 8.0; 107.6, 25.0; 117.6, 35.0; 122.6, 75.0; 125.6, 95.0; 131.6, 95.0; 132.6, 50.0; 134.0, 50.0; 134.6, 95.0; 140.6, 95.0; 142.6, 1.0; 190.0, 1.0. For data-independent acquisition (DIA), master MS scans were acquired over an m/z range of 400–900 at a resolution of 60,000, with an automatic gain control (AGC) target of 1E6 charges and a maximum ion injection time of 128 ms. Precursor ions were fragmented by higher-energy collisional dissociation (HCD) using a normalized collision energy of 30. MS/MS spectra were acquired at a resolution of 30,000 over an m/z range of 400–900, using a 10 Da isolation window, an AGC target of 1E6, and a maximum ion injection time of 64 ms. MSX isochronous ion injection times were enabled, and the MSX count was set to 1. The DIA data for the peptides derived from UCI female line was acquired on Orbitrap Exploris 480 coupled with a Vanquish Neo UHPLC system using comparable settings. The proteomics data is available on MassIVE repository (https://massive.ucsd.edu/) with identifier MSV000101439.

### Data processing and functional enrichment

The Thermo .raw files were converted to .mzML format using MSConvert (v3.0.24239.0) with peak picking enabled. The resulting .mzML files were analyzed in FragPipe (v23.0), incorporating MSFragger (v4.2) and DIA-NN (v2.1.0), using the DIA_SpecLib_Quant workflow. Database searches were performed against the Homo sapiens UniProt/Swiss-Prot reference proteome (Proteome ID: UP000005640; 20,417 entries), supplemented with common contaminants and decoy sequences. Search parameters were as follows: strict trypsin specificity with up to 4 missed cleavages; precursor mass tolerance of ±10 ppm; fragment mass tolerance of 20 ppm; variable modifications including protein N-terminal acetylation and methionine oxidation; fixed carbamidomethylation of cysteine residues; and a false discovery rate (FDR) threshold of 1%. Further data processing and statistical analysis were performed on the protein-level output file from FragPipe (pg_matrix.tsv) with RomicsProcessor v1.1.0 (https://github.com/PNNL-Comp-Mass-Spec/RomicsProcessor). Briefly, the LFQ intensities were log2-transformed, filtered to allow maximal missingness of 40% within each group, and normalized by median. The missing values remaining post-filtering (<0.5%) were imputed following a previously described method by Tyanova et al. An unpaired t-test was performed to compare the groups corrected with Benjamini-Hochberg. Finally, a functional enrichment analysis was performed on the differentially upregulated and differentially downregulated (p-adj < 0.05, |Log2FC cutoff| > 0.5) proteins using Metascape^97^.

### Bulk RNA sequencing of isogenic iMG

For GW3965 treated conditions, iMG were exposed to 10 μM GW3965 (Apexbio Tech, A8444-5mg) or DMSO for 24 h before RNA isolation. iMG RNA was isolated from the cell pellets using the Direct-zol RNA Microprep (Zymo, R2061). Novogene services were used for RNA library preparation and sequencing. Sequencing was carried out using an Illumina NovaSeq X Plus Series (PE150). Reads were mapped to the human hg38 reference genome using STAR (v. 2.7.11b). Raw read counts were generated with Salmon (v. 1.10.3). Differential expression was performed using the R package DESeq2. Pathway analysis was done using Metascape^97^.

### ROSMAP microglia analysis

The ROSMAP snRNA-seq data were processed, analyzed and annotated by Mathys et al^89^. Within the immune cells, there are 3 microglia clusters that were then used in this study for subsequent differential analysis across different APOE subtypes. There are only 8 APOE44 donors in the ROSMAP cohort, with at least 10 microglia cells each. In contrast, there are 246 APOE33 donors. For this reason, we decided to randomly down-sample the APOE33 group to 8 samples to perform the differential analysis using pseudobulk profiles, followed by DESeq2, between APOE44 and APOE33. Pathway analysis was done using Metascape. All donors with at least 10 microglia cells and having a diagnosis of AD or CTRL were used to compare the distribution of BHLHE40-/41- cells, CDKN2A+ (p16+) cells, and ALOX5AP+ cells across APOE subtypes, using chi-squared test to evaluate significance.

